# Cell line-specific network models of ER+ breast cancer identify PI3Kα inhibitor sensitivity factors and drug combinations

**DOI:** 10.1101/2020.08.25.261370

**Authors:** Jorge Gómez Tejeda Zañudo, Pingping Mao, Clara Alcon, Kailey J. Kowalski, Gabriela N. Johnson, Guotai Xu, José Baselga, Maurizio Scaltriti, Anthony Letai, Joan Montero, Reka Albert, Nikhil Wagle

## Abstract

Durable control of invasive solid tumors necessitates identifying therapeutic resistance mechanisms and effective drug combinations. A promising approach to tackle the cancer drug resistance problem is to build mechanistic mathematical models of the signaling network of cancer cells, and explicitly model the dynamics of information flow through this network under distinct genetic conditions and in response to perturbations.

In this work, we used a network-based mathematical model to identify sensitivity factors and drug combinations for the PI3Kα inhibitor alpelisib, which was recently approved for ER+ *PIK3CA* mutant breast cancer. We experimentally validated the model-predicted efficacious combination of alpelisib and BH3 mimetics (e.g. MCL1 inhibitors) in ER+ breast cancer cell lines. We also experimentally validated the reduced sensitivity to alpelisib caused by FOXO3 knockdown, which is a novel potential resistance mechanism. Our experimental results showed cell line-specific sensitivity to the combination of alpelisib and BH3 mimetics, which was driven by the choice of BH3 mimetics. We find that cell lines were sensitive to the addition of either MCL1 inhibitor s63845 alone or in combination with BCL-XL/BCL-2 inhibitor navitoclax, and that the need for the combination of both BH3 mimetics was predicted by the expression of BCL-XL. Based on these results, we developed cell line-specific network models that are able to recapitulate the observed differential response to alpelisib and BH3 mimetics, and also incorporate the most recent knowledge on resistance and response to PI3Kα inhibitors.

Overall, we present an approach for the development, experimental testing, and refining of mathematical models, which we apply to the context of PI3Kα inhibitor drug resistance in breast cancer. Our approach predicted and validated PI3Kα inhibitor sensitivity factors (FOXO3 knockdown) and drug combinations (BH3 mimetics), and illustrates that network-based mathematical models can contribute to overcoming the challenge of cancer drug resistance.

## Introduction

Overcoming intrinsic or acquired resistance to cancer drug therapies is one the biggest challenges faced today by patients, oncologists, and cancer researchers (Vasan, Baselga, and Hyman 2019; Konieczkowski, Johannessen, and Garraway 2018). This is particularly the case in the setting of metastatic disease, which accounts for the majority of cancer deaths (Steeg 2016; Weigelt, Peterse, and van ‘t Veer 2005). The challenge underlying cancer drug resistance is exemplified by metastatic estrogen receptor positive (ER+) breast cancer, the most common type of metastatic breast cancer. Despite the availability of multiple effective drug therapies (Waks and Winer 2019) (endocrine and targeted therapies, CDK4/6 inhibitors, among others), ER+ metastatic breast cancer eventually develops resistance to all available treatments. As a result, long-term patient survival has not greatly improved, with a median prognosis of around 3-4 years after the development of metastatic disease (Caswell-Jin et al. 2018; Waks and Winer 2019).

One of the most recently approved drugs to treat ER+ breast cancer is alpelisib (previously known as BYL719), which is a PI3Kα-specific inhibitor approved for use in combination with fulvestrant (a selective estrogen receptor degrader, a type of ER inhibitor) for ER+ metastatic breast cancer with activating *PIK3CA* mutations (André et al. 2019). Multiple lines of clinical evidence support PTEN loss-of-function as a clinically relevant resistance mechanism to alpelisib and other PI3Kα inhibitors (Juric et al. 2015; Costa et al. 2020; Razavi et al. 2020). There are several other potential PI3Kα inhibitor resistance mechanisms that have been identified through *in vitro* or *in vivo* studies (Elkabets et al. 2013; Le et al. 2016; Castel et al. 2016; Costa et al. 2015; Schwartz et al. 2015; Bosch et al. 2015; Kodack et al. 2017; Donnella et al. 2018; Hopkins et al. 2018), but it is still an open question if these will drive resistance in patients and if they will be common enough to be clinically relevant. It is also likely that there are currently unidentified PI3Kα resistance mechanisms that will be found as the landscape of intrinsic and acquired resistance mechanisms is revealed, as has happened in the case of endocrine therapies (Razavi et al. 2018; Bertucci et al. 2019; Nayar et al. 2019; Mao et al. 2020) and CDK4/6 inhibitors (Wander et al. 2020; Li et al. 2018).

How can we identify these currently unknown PI3Kα resistance mechanisms? Based on what we have learned from other targeted drugs (e.g. BRAF(Lito, Rosen, and Solit 2013) or EGFR inhibitors(Sun and Bernards 2014)), resistance mechanisms act by reactivating the targeted pathway or by activating alternative oncogenic pathways that bypass it (Vasan, Baselga, and Hyman 2019; Konieczkowski, Johannessen, and Garraway 2018; Wagle et al. 2011). Therefore, a promising avenue to identify such resistance mechanisms is to use approaches that combine a deep understanding of the oncogenic pathways driving ER+ *PIK3CA* mutant breast cancer, their cross-talk, and their integration by cancer cells to make the cellular decisions that result in tumor growth. One such approach is provided by mathematical models of cancer (Clarke and Fisher 2020; Rockne et al. 2019), particularly dynamic network models, which mechanistically model the dynamics of the molecular network underlying cancer cells (G. T. Zañudo, Steinway, and Albert 2018; Saez-Rodriguez and Blüthgen 2020).

Dynamic network models are based on mechanistic knowledge and explicitly model how information flows through signaling networks under distinct genetic conditions (e.g. in a context where known or potential resistance mechanisms are active) and in response to perturbations (e.g. drugs or environmental signals). A particularly attractive property of dynamic network models in the context of cancer is that these models can be customized to become specific to particular cancer cell lines, or even to individual patient tumors, by introducing known genetic alterations and/or by mimicking the known response (or lack thereof) to a perturbation (Saez-Rodriguez and Blüthgen 2020). Tested and validated network models can predict the degree of effectiveness of single-agent and combinatorial drug interventions, and predict potential mechanisms of drug resistance, which can then be used to prioritize drug combinations for *in vitro* and *in vivo* testing in relevant cancer model systems (G. T. Zañudo, Steinway, and Albert 2018; Saez-Rodriguez and Blüthgen 2020; Gómez Tejeda Zañudo, Scaltriti, and Albert 2017; Steinway et al. 2015; Béal et al. 2018; Fröhlich et al. 2018; Barrette, Bouhaddou, and Birtwistle 2018; Fey et al. 2015).

We previously built a network-based mathematical model of oncogenic signal transduction in ER+ *PIK3CA* mutant breast cancer (Fig. 1A) (Gómez Tejeda Zañudo, Scaltriti, and Albert 2017). The model integrated the mechanistic knowledge of the response/resistance to PI3Kα inhibitors and other drugs relevant in the context of ER+ breast cancer (e.g. CDK4/6 inhibitors, ER inhibitors, or mTORC1 inhibitors). The model was based on the combined knowledge of *in vitro, in vivo*, and clinical literature; it recapitulated the drug resistance phenotype of a variety of resistance mechanisms to PI3Kα inhibitors (e.g. PTEN loss-of-function, mTORC1 activity, and PIM overexpression) and other drugs (e.g. RB1 loss in response to CDK4/6 inhibitors, and ER transcriptional reactivation in response to ER inhibitors). By exhaustively testing the effect of all single- and double-element perturbations, the model predicted potential resistance mechanisms and efficacious drug combinations for PI3Kα inhibitors (Fig. 1B). In particular, it predicted the knockdown of tumor suppressors FOXO3, RB1, or p21 and p27 as potential PI3Kα inhibitor resistance mechanisms, and identified BH3 mimetics targeting anti-apoptotic proteins MCL1, BCL-XL, and BCL2 as efficacious drug combinations.

**Fig 1.**
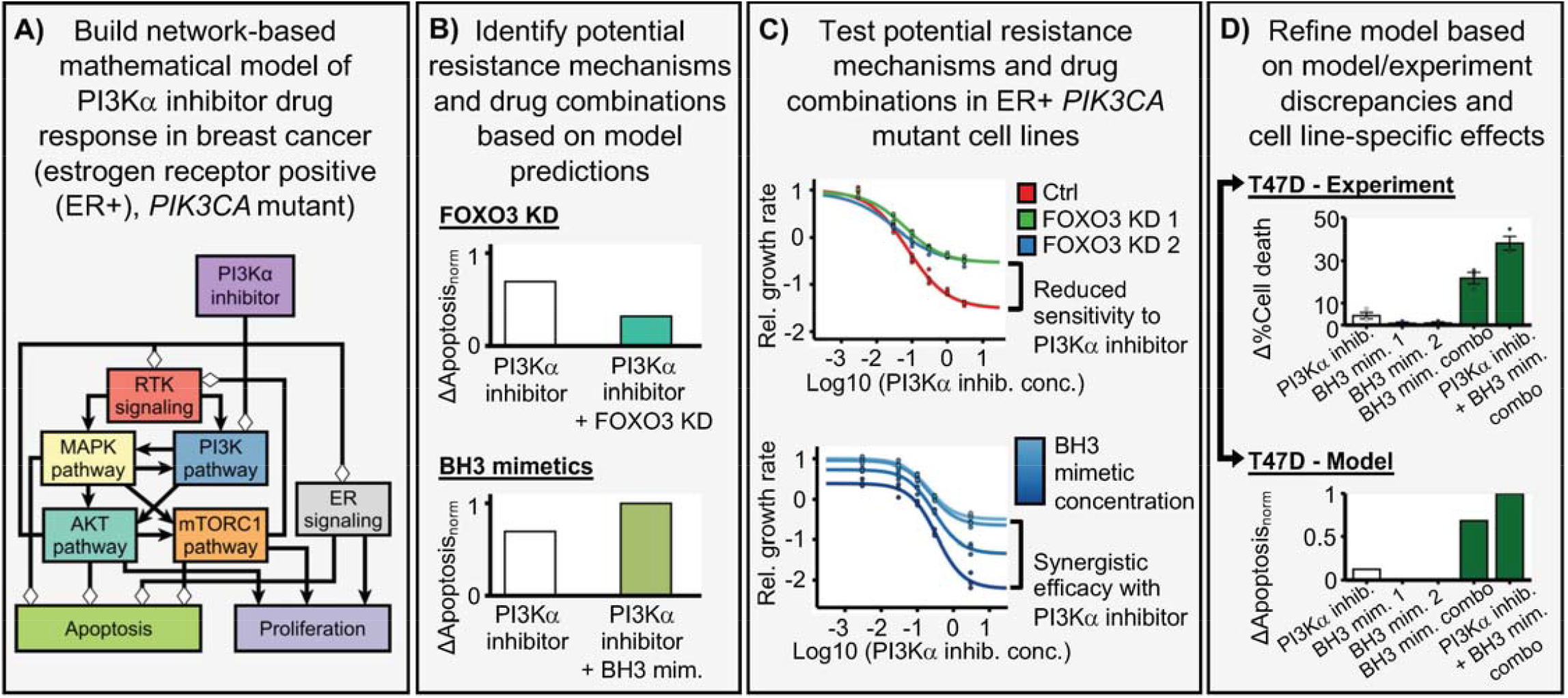
Outline of the research strategy followed in this work. (A) We use a network-based mathematical model of oncogenic signal transduction in estrogen receptor positive (ER+), *PIK3CA* mutant breast cancer. This model integrates the mechanistic knowledge of the response/resistance to PI3Kα inhibitors and recapitulates the drug resistance phenotype of a variety of known resistance mechanisms. (B) Based on this model, we predict potential resistance mechanisms (e.g. FOXO3 knockdown) and drug combinations (BH3 mimetics) for PI3Kα inhibitors. (C) We experimentally test these predicted resistance mechanisms and drug combinations in ER+, *PIK3CA* mutant breast cancer cell lines. Based on these experiments, we can verify the model’s predictions and pinpoint any discrepancies between the model and the experimental results. (D) We use the experimental results and the discrepancies between the model and the experiments to refine the network-based mathematical model and incorporate the observed cell line-specific effects in the model.

In this work we experimentally test the above predictions of our mathematical model (Fig. 1C). We find that some of the model’s predictions are verified experimentally, particularly the efficacy of the drug combination of alpelisib and BH3 mimetics, and the reduced sensitivity to PI3Kα inhibitors caused by FOXO3 knockdown. We also find some discrepancies between our experimental findings and the model, some of which are cell line-specific. In particular we find a cell line-specific response to BH3 mimetics in MCF7 and T47D, two of the most commonly used cell line models for ER+ *PIK3CA* mutant breast cancer. Finally, we construct cell line-specific models that incorporate and recapitulate the most recent work on PI3Kα inhibitors (Hopkins et al. 2018; Donnella et al. 2018), including our experimental results. These experimental results—and in particular the discrepancies between the model and the experiments—are then used to refine the network-based mathematical model and incorporate the observed cell line-specific effects into the model (Fig. 1D).

## Results

### PI3Kα inhibitors and BH3 mimetics are a synergistically efficacious drug combination and induce apoptosis in ER+ *PIK3CA*-mutant breast cancer cell lines

In the network model, PI3Kα inhibition has a cytotoxic effect mediated by the increased activity of the pro-apoptotic BCL-2 family members BIM and BAD (through upregulation of FOXO3-mediated transcription (Sunters et al. 2003) and through a decrease of an inactivating AKT-mediated phosphorylation (She et al. 2005), respectively). The cytotoxic effect of PI3Kα inhibition is countered by the presence of anti-apoptotic BCL-2 family members MCL1 and BCL2 (which in the model represents BCL-2 and other anti-apoptotic family members such as BCL-XL). As a result, the combined inhibition of PI3Kα and anti-apoptotic family members MCL1, BCL-2, or BCL-XL, each of which can be targeted using BH3 mimetics, is predicted to be an effective and synergistic drug combination (Gómez Tejeda Zañudo, Scaltriti, and Albert 2017). Specifically, the model predicts that inhibition of the anti-apoptotic family members would be more effective to induce apoptosis than the combinations used with alpelisib in clinical practice (fulvestrant) and in clincal trials (palbociclib, everolimus), all of which predominantly inhibit cell proliferation.

To test the effect of combining PI3Kα inhibitors and BH3 mimetics of anti-apoptotic proteins, we first treated MCF7 cells (ER+, *PIK3CA*-mutant) with PI3Kα inhibitor alpelisib and MCL1 inhibitor s63845 (a BH3 mimetic (Kotschy et al. 2016)) separately and in combination using both a long-term colony formation assay (2 weeks, Fig. 2A) and a short-term cell viability assay (6 days, Fig. 2B). In both cases the combination of alpelisib and s63485 was more efficacious than each drug on its own. We calculated the synergy score of the combination using MuSyC (Wooten et al. 2019; Meyer et al. 2019) and found that it was synergistically efficacious (*β*_*obs*_ *= 0*.*45*, where *β*_*obs*_ *>0* means the combination is synergistically efficacious and *β*_*obs,null*_ *= 0*.*08* is the null-hypothesis value expected from doubling the alpelisib concentration). To compare the synergistic efficacy of the alpelisib + s63845 combination with other proposed alpelisib drug combinations, we performed drug response curves of the combination of alpelisib with fulvestrant (the FDA-approved combination for the treatment for ER+ *PIK3CA*-mutant metastatic breast cancer (André et al. 2019; Bosch et al. 2015)), palbociclib (Vora et al. 2014), or everolimus (which had previously been found to increase alpelisib-induced cell death (Elkabets et al. 2013)), and calculated their synergy scores (Fig. 2C, Suppl. Fig. S1). We found that all the tested drug combinations were synergistically efficacious (i.e. *β*_*obs*_ *> 0*, with their range being 0.24 - 0.45; Fig. 2C), but the alpelisib + s63845 combination had the highest synergy score among all the combinations (*β*_*obs*_ *= 0*.*45*).

**Fig 2.**
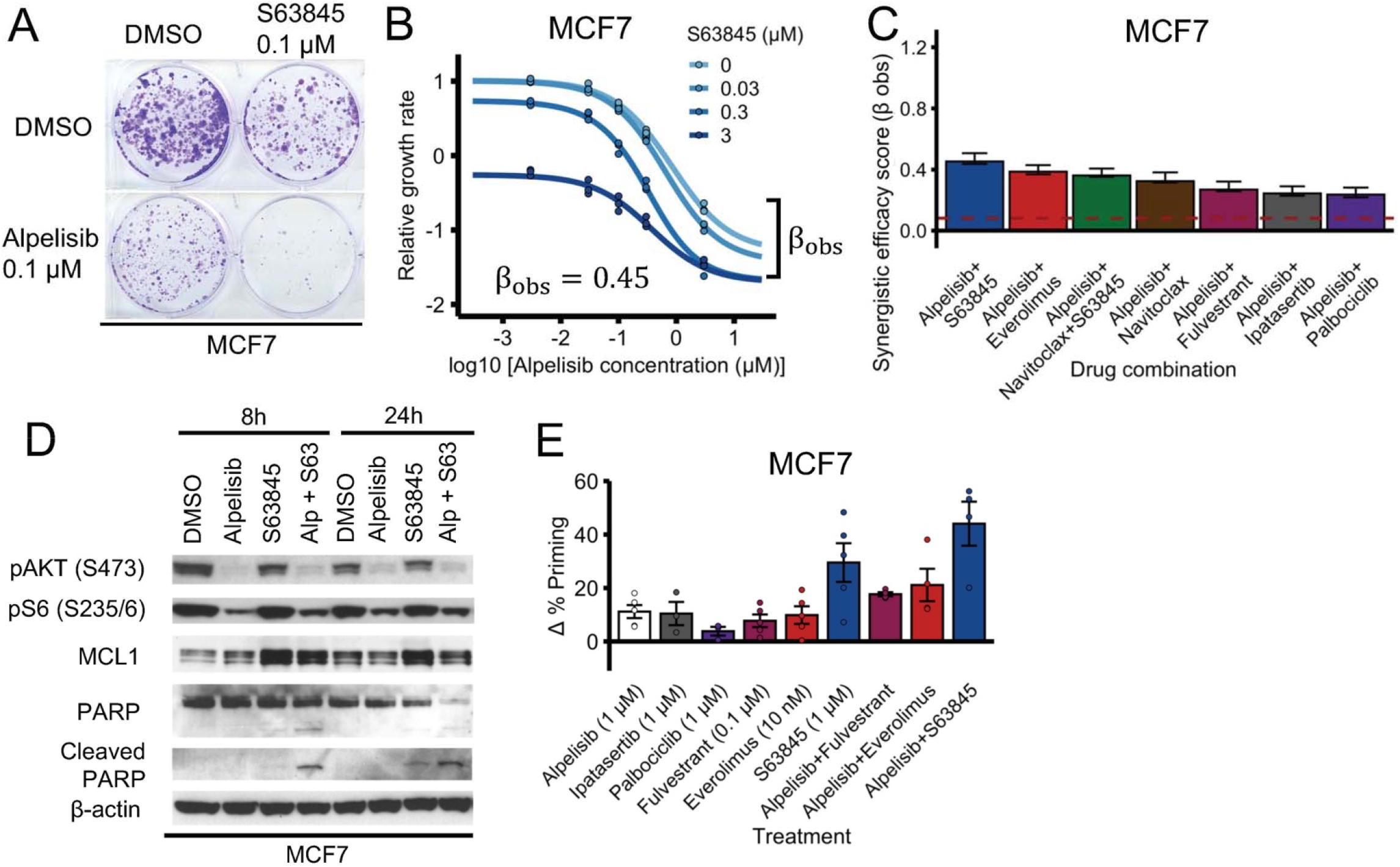
PI3Kα inhibitors and MCL1 inhibitors are a synergistically efficacious combination and induce apoptosis in MCF7. (A, B) PI3Kα inhibitor alpelisib and MCL1 inhibitor s63845 is an effective and synergistically efficacious drug combination. MCF7 cells were treated with alpelisib, s63845, and their combination. For panel A, cells a colony formation assay was performed after 18 days of drug treatment. For panel B, CellTiter-Glo assay was performed before drug treatment and after 6 days of drug treatment to measure cell viability. The relative growth rate is calculated with respect to the DMSO control treatment using cell viability. Efficacy scores *β*_*Obs*_ are such that *β*_*Obs*_ > 0 means synergistic behavior and *β*_*Obs*_ < 0 means antagonistic behavior, and are calculated using MuSyC (Meyer et al. 2019). (C) The combination of alpelisib and s63845 is equally synergistically efficacious (or more) than clinically relevant drugs like fulvestrant, everolimus, and palbociclib. The dotted red line (*β*_*Obs*_ = 0.08) is the null-hypothesis synergy score value expected from doubling the alpelisib combination (i.e., a combination of alpelisib with itself). Experiments were done and synergy scores *β*_*Obs*_ were calculated as in panel B. (D) Combined alpelisib and s63845 result in an increase in apoptosis (as measured by cleaved PARP levels) compared to each separate drug. Consistent with the known effects of alpelisib and s63845, we observe that alpelisib (0.1 μM) decreased phosphorylation of AKT (S473) and S6 (S235/6) (downstream targets of PI3K signaling), and that s63845 (0.1 μM) increased MCL1 protein levels (due to s63845’s binding to MCL1, which results in its inhibition (Kotschy et al. 2016)). (E) The combination of alpelisib with s63845 results in an increased apoptosis priming compared to the combination with clinically relevant drugs like fulvestrant and everolimus. Δ% apoptosis priming for a treatment was measured using dynamic BH3 profiling by comparing treated vs untreated cells after 16 h under increased concentration of BIM peptide.

To determine if the effect of the alpelisib + s63845 combination is mediated by apoptosis or other mechanisms, we treated MCF7 with each drug separately and in combination, and measured the levels of cleaved PARP and total PARP (Fig. 2D). We found that alpelisib + s63845 induced increased levels of cleaved PARP compared to each drug alone at both time points (8 and 24 h), with the effect being more evident at the early (8 h) time point (Fig. 2D). To contextualize the apoptotic effect of the alpelisib + s63845 combination, we tested the effect of single agent and drug combinations and measured the resulting apoptotic priming using dynamic BH3 profiling (Montero et al. 2015) (Fig. 2E). The alpelisib + s63845 combination showed the highest increase in *Δ*% priming (which measures apoptotic sensitivity) of all combinations, and this effect was larger than each drug on its own. In particular, the alpelisib + s63845 combination showed a higher *Δ*% priming than alpelisib + fulvestrant and alpelisib + everolimus.

We also performed similar experiments in T47D cells, another ER+ breast cancer cell line. In contrast to MCF7, T47D cells did not show a strong synergistic effect under the combination of alpelisib + s63845 (Fig. 3B, Suppl. Fig S2A). However, the addition of navitoclax (a BH3 mimetic targeting BCL-XL and BCL-2) to the combination of alpelisib and s63845 recapitulated the effects seen in the MCF7 cells (Fig. 3A-B). The alpelisib + s63845 + navitoclax combination had a strong synergistically efficacious effect (*β*_*obs*_ *= 1*.*10*; Fig. 3A) and had a significantly higher synergy score than other combinations tested (Fig. 3B). The combination of alpelisib + s63845 + navitoclax in T47D also showed increased apoptosis, measured by cleaved PARP and PARP, which is consistent with the results in MCF7 cells (Fig. 3D). The increased apoptosis was further confirmed by measuring cell death through Annexin V staining (Fig. 3C).

**Fig 3.**
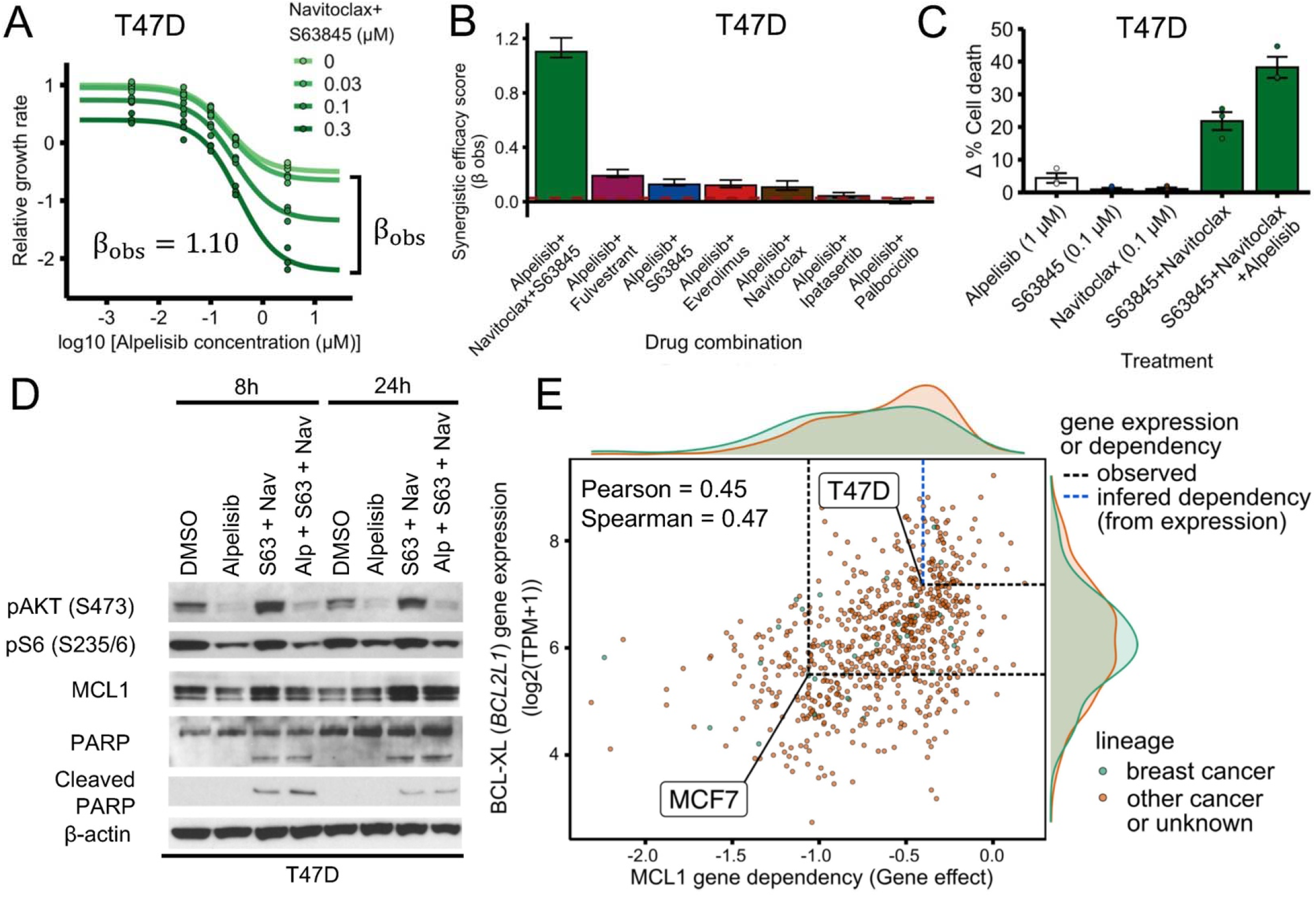
PI3Kα inhibitors and BH3 mimetics are a synergistically efficacious combination and induce apoptosis in T47D. (A) PI3Kα inhibitor alpelisib, MCL1 inhibitor s63845, and BCL-2/BCL-XL inhibitor navitoclax is an effective and synergistically efficacious drug combination. T47D cells were treated with alpelisib, s63845 and navitoclax (1 μM), and their combination. CellTiter-Glo assay was performed before drug treatment and after 6 days of drug treatment to measure cell viability. The relative growth rate is calculated with respect to the DMSO control treatment using cell viability. Efficacy scores *β*_*obs*_ are such that *β*_*obs*_ > 0 means synergistic behavior and *β*_*obs*_ < 0 means antagonistic behavior, and are calculated using MuSyC (Meyer et al. 2019). (B) The combination of alpelisib, s63845, and navitoclax is more synergistically efficacious than clinically relevant drugs like fulvestrant, everolimus, and palbociclib. The dotted red line (*β*_*obs*_ = 0.03) is the null-hypothesis synergy score value expected from doubling the alpelisib combination (i.e., a combination of alpelisib with itself). Experiments were done and synergy scores *β*_*obs*_ were calculated as in Fig. 2. (C, D) Combined alpelisib, s63845, and navitoclax result in an increase in apoptosis compared to each separate drug. Apoptosis was measured using annexin V (panel C) and cleaved PARP levels (panel D). Δ% cell death for a treatment was measured using annexin V by comparing treated vs DMSO control cells after 72 h (panel C). The decreased phosphorylation of AKT (S473) and S6 (S235/6), and increased levels of MCL1 are consistent with the known effects of alpelisib and s63845 (see Fig. 2). Drug concentrations used were: 0.1 μM for alpelisib, 0.3 μM for s63845, and 0.3 μM for navitoclax (panel D). (E) Sensitivity to MCL1 knockout in CRISPR/Cas-9 loss-of-function screens (DepMap) is strongly correlated with BCL-XL gene expression. Based on T47D’s BCL-XL gene expression, T47D is inferred to be less sensitive to MCL1 knockout than MCF7, consistent with T47D requiring navitoclax to be sensitive to combined alpelisib and s63845. Sensitivity to gene knockout is measured using gene effect, which is such that gene effect=0 means no effect and gene effect=-1 is the median effect of known common essential genes. Inferred gene effect is shown with a dashed blue line; gene expression or gene effect from the dataset is shown with a dashed black line.

Given that the addition of BCL-XL/BCL-2 inhibitor navitoclax sensitized T47D cells to the combination of alpelisib + s63845, we reasoned that this was because T47D cells had a higher expression of BCL-XL and/or BCL-2 compared to MCF7 cells. To test this hypothesis, we looked at the gene and protein expression of MCF7 and T47D cells in the Cancer Cell Line Encyclopedia (CCLE) (Nusinow et al. 2020; Ghandi et al. 2019). Gene and protein expression of BCL-XL was higher in T47D compared to MCF7, consistent with our hypothesis (Supp. Fig S2B;|*Δ log*_2_ *(TPM + 1)*| *= 1*.*68*, difference robust z-score |*z*| *= 1*.*08*; |*Δ Relative protein expression*| = 0.64, difference robust z-score |*z*| *= 0*.*59*). For the case of BCL-2, there was little difference in protein expression and the difference in gene expression was in the opposite direction of the effect we are trying to explain (higher gene expression in MCF7 compared to T47D vs increased sensitivity in T47D when adding navitoclax) (Supp. Fig S5B;|*Δ log*^*2*^ *(TPM + 1)*| *= 4*.*14*, difference robust z-score |*z*| *= 2*.*19*;|*Δ Relative protein expression*| = 0.13, difference robust z-score |*z*| *= 0*.*09*).

To further test whether BCL-XL expression can serve as a marker for cells that require inhibition of BCL-XL to become sensitive to MCL1 inhibitors, we looked at how the effect of MCL1 gene knockout correlated with BCL-XL gene expression in various cell lines using the CRISPR/Cas9 loss-of-function genetic screens of the Cancer Dependency Map (DepMap) (Meyers et al. 2017; Dempster et al. 2019). Consistent with the hypothesis of BCL-XL as a marker, sensitivity to MCL1 gene knockout (MCL1 gene dependency) was strongly correlated with reduced BCL-XL expression (Fig. 3E; *Pearson correlation = 0*.*45, *p* = 1*.*9 x 10*^*-37*^; *Spearman correlation = 0*.*47, *p* = 7*.*2 x 10*^*-41*^) and this was the strongest gene expression correlate (Pearson) among all genes. There was no strong correlation between sensitivity to MCL1 gene knockout and MCL1 expression (*Pearson correlation = −0*.*03, *p* = 4*.*3 x 10*^*-1*^; *Spearman correlation = −0*.*03, *p* = 3*.*6 x 10*^*-1*^) or BCL-2 expression (*Pearson correlation = −0*.*15, *p* = 4*.*3 x 10*^*-*^ ; *Spearman correlation = −0*.*10, *p* = 8*.*4 x 10*^*-3*^). Consistent with our experimental findings, MCF7 was in the low end of BCL-XL expression (Fig. 3E; *log*^*2*^ *(TPM + 1) = 5*.*50*, robust z-score *z = −0*.*62*) and was sensitive to MCL1 gene knockout (Fig. 3E; *gene effect = −1*.*06*, where a *gene effect* of 0 means no effect, while a *gene effect* of −1 means the same effect as an essential gene). T47D was in the high end of BCL-XL expression (Fig. 3E; *log*_*2*_ *(TPM + 1) = 7*.*19*, robust z-score *z = 0*.*94*) and, although it is not in the DepMap database, the inferred sensitivity to MCL1 gene knockout was less than half of that for MCF7 (Fig. 3E; *median(gene effect) = −0*.*40* for the 57 cell lines within *±2*.*5%* of the *log*_*2*_ *(TPM + 1)* of T47D’s MCL1 gene expression). We also found consistent results for the effect of MCL1 gene knockout and BCL-XL expression when looking at other publicly available loss-of-function (RNAi) screen datasets (Supp. Fig. S2C), although the dynamic range of gene effects for MCL1 was much smaller for these RNAi screen datasets.

In summary, these results support the predictions of the model that (1) the combined inhibition of PI3Kα and the relevant anti-apoptotic family members (MCL1 for MCF7; MCL1 and BCL-XL for T47D) is an effective and synergistic drug combination, and (2) combined alpelisib and BH3 mimetics have a larger cell death effect than each drug alone and other effective drug combinations (alpelisib + everolimus, alpelisib + fulvestrant).

### FOXO3 knockdown is a potential resistance mechanism to PI3Kα inhibitors

In the network model, PI3Kα inhibition increases FOXO3 nuclear localization and transcriptional activity through AKT-mediated dephosphorylation of FOXO3 (Brunet et al. 1999; Castel et al. 2016; Muranen et al. 2012; Chandarlapaty et al. 2011). Increased FOXO3 nuclear activity upregulates the expression of downstream targets that result in the suppression of cell growth and induction of cell death (Hu et al. 2004; Brunet et al. 1999; Medema et al. 2000; Castel et al. 2016). As a result, the network model predicts that knockdown of FOXO3 is a potential resistance mechanism to PI3Kα inhibitors and that this reduced sensitivity is a result of blocking FOXO3 transcriptional activity (Gómez Tejeda Zañudo, Scaltriti, and Albert 2017).

To test this prediction, we generated MCF7 and T47D cell lines with FOXO3 knockdown (KD) using CRISPR-Cas9 technology and verified the knockdown of FOXO3 by western blot (Fig. 4A). We tested the response of FOXO3 KD MCF7 and T47D cells to two PI3Kα inhibitors (alpelisib and taselisib) and found that both cell lines show reduced sensitivity to both PI3Kα inhibitors when compared to control cells with a non-targeting (NT) sgRNA (Fig. 4B; two-sided t-test *p < 0*.*01* for each sgRNA vs NT control comparison, fold change 1.4 - 2.2 compared to NT control for MCF7, fold change 1.4 - 1.5 compared to NT control for T47D). To further test this reduced sensitivity to PI3Kα inhibitors, we performed drug response curves to varying concentrations of alpelisib or taselisib (Fig. 4C-D). We found that the drug response curves show a reduced sensitivity to PI3Kα inhibitors in FOXO3 KD MCF7 cells as compared to control cells, (Fig. 4C), consistent with the resistant phenotype in Fig. 4B. For FOXO3 KD T47D cells we did not see significant effect on PI3Kα inhibitor sensitivity from FOXO3 knockdown in the drug response curves (compare Fig. 4D and 3B), a result that may be attributed to the methodology being insufficiently sensitive to measure moderate effects. (Supp. Fig. S3).

**Fig 4.**
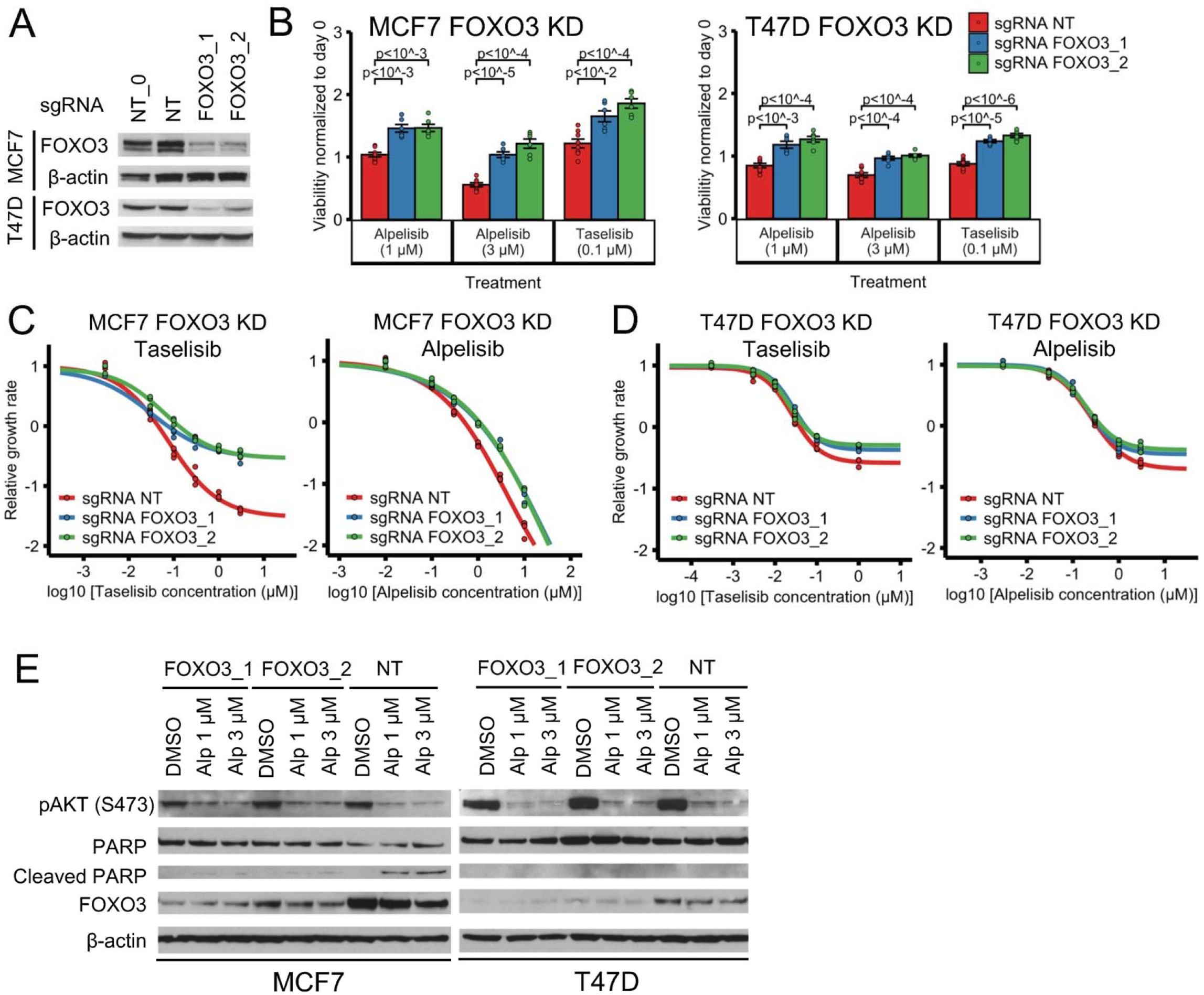
FOXO3 knockdown reduces sensitivity to PI3Kα inhibitors in ER+ breast cancer cell line models. (A) Stable cell lines generated using CRISPR FOXO3 knockdown have reduced proteins levels of FOXO3. NT and NT_0 are two distinct non-targeting control guides; FOXO3_1 and FOXO3_2 are two distinct guides targeting FOXO3. (B-D) FOXO3 knockdown reduces sensitivity to PI3Kα inhibitors alpelisib and taselisib in ER+ breast cancer cell lines MCF7 and T47D. CellTiter-Glo assay was used to measure cell viability before drug treatment and after 6 days of drug treatment. An unpaired, two-sided t-test was used for comparison between groups (panel B). Experiments in panel B single-dose experiments were optimized for the tested PI3Kα inhibitor concentrations: panel B used a larger number of initial cells than panels C and D (5,000 vs 1,000 cells), and measure the viability with respect to the cells before treatment. Panels C and D calculate the relative growth rate with respect to DMSO control. (E) FOXO3 knockdown reduces PI3Kα-inhibitor-induced cell death in MCF7. Cell death was measured using cleaved PARP levels. The decreased phosphorylation of AKT (S473) is consistent with the known downregulation of PI3K signaling by alpelisib.

Many drug resistance mechanisms for targeted therapies involve reactivation of the signaling pathway inhibited by targeted therapy. However, according to the model, FOXO3 KD does not reactivate the PI3K pathway. We decided to test this experimentally by measuring AKT phosphorylation, which is indicative of PI3K pathway activation. We found that that AKT(S473) phosphorylation was similar when comparing FOXO3 KD and control cells at baseline and in response to alpelisib in both MCF7 and T47D (Fig. 4E). We conclude that FOXO3 KD does not reactivate PI3K pathway activity. This is consistent with the current knowledge on FOXO3, which indicates that it mainly acts downstream of the PI3K pathway but might also act upstream of the PI3K pathway due to feedback activation, and suggests that the effect of feedback activation by FOXO3 is weak in this context.

One important prediction of the model is that FOXO3 KD reduces PI3Kα inhibitor sensitivity by decreasing drug-induced apoptosis. To test this, we determined the effect of alpelisib on cell death by measuring the levels of cleaved PARP and PARP in FOXO3 KD vs control cells (Fig. 4E). We found that the FOXO3 KD MCF7 cells have reduced levels of cleaved PARP compared to control (Fig. 4E, left). We did not see this difference in cleaved PARP for T47D (Fig. 4E, right). The reduced levels of cleaved PARP seen only in MCF7 FOXO3 KD cells is consistent with the reduced cytotoxic effect seen in the alpelisib drug response curves of FOXO3 KD in MCF7 but not in T47D (Fig. 4C-D).

Overall, we found that FOXO3 KD reduces sensitivity to PI3Kα inhibitors in both T47D and MCF7, and that this effect was more pronounced in MCF7 cells, which is mediated by a reduction in drug-induced apoptosis in MCF7. These results are consistent with the predictions of the model and suggest that downregulation of FOXO3 is a potential resistance mechanism for PI3Kα inhibitors.

### CDKN1B (p27) knockdown does not have a strong effect on PI3Kα inhibitor sensitivity

The network model contains a node, *p21/p27*, that merges the tumor suppressor CDKN1B (p27) and its paralogue CDKN1A (p21). In the model, PI3Kα inhibition results in an increased activity of this node, mediated by the upregulation of FOXO3-mediated *CDKN1B* transcription (Medema et al. 2000) and a decrease in AKT-mediated p27 and p21 phosphorylation (Viglietto et al. 2002; Liang et al. 2002; Zhou et al. 2001). p27 and p21 exert their tumor suppressor function by binding and inactivating the cyclin E-CDK2 complex (Sherr and Roberts 1999; Polyak et al. 1994; Wander, Zhao, and Slingerland 2011; Planas-Silva and Weinberg 1997), which upon release from p27 or p21 promotes the G1-S transition and progression into the cell cycle. As a consequence, the model predicts that inactivation of the merged *p21/p27* node will interfere with the cytostatic effect of PI3Kα inhibition, and result in a reduced sensitivity to PI3Kα inhibitors (Gómez Tejeda Zañudo, Scaltriti, and Albert 2017). As the model does not specify the contribution of p27 and p21 to this merged node, inactivation of this merged node might require simultaneous knockdown of CDKN1B and CDKN1A, or, if the contribution of p21 is minor, might be reproduced by knockdown of CDKN1B only. Given that p27 but not p21 has been shown to be transcriptionally upregulated by AKT-mediated FOXO3 activation (Medema et al. 2000; X. Zhang et al. 2011), we chose to investigate the contribution of p27 by knocking down CDKN1B only.

We generated MCF7 and T47D cell lines with CDK1NB knockdown, which we verified through western blotting (Fig. 5A). We then tested the response of CDKN1B KD cells to the PI3Kα inhibitors alpelisib and taselisib (Fig. 5B-D). We found that the CDKN1B KD cell lines did not show a consistent reduced sensitivity to PI3Kα inhibition compared to control, neither in the drug response curves nor in the further optimized drug treatment experiment at selected doses. (Fig. 54B-D).

**Fig 5.**
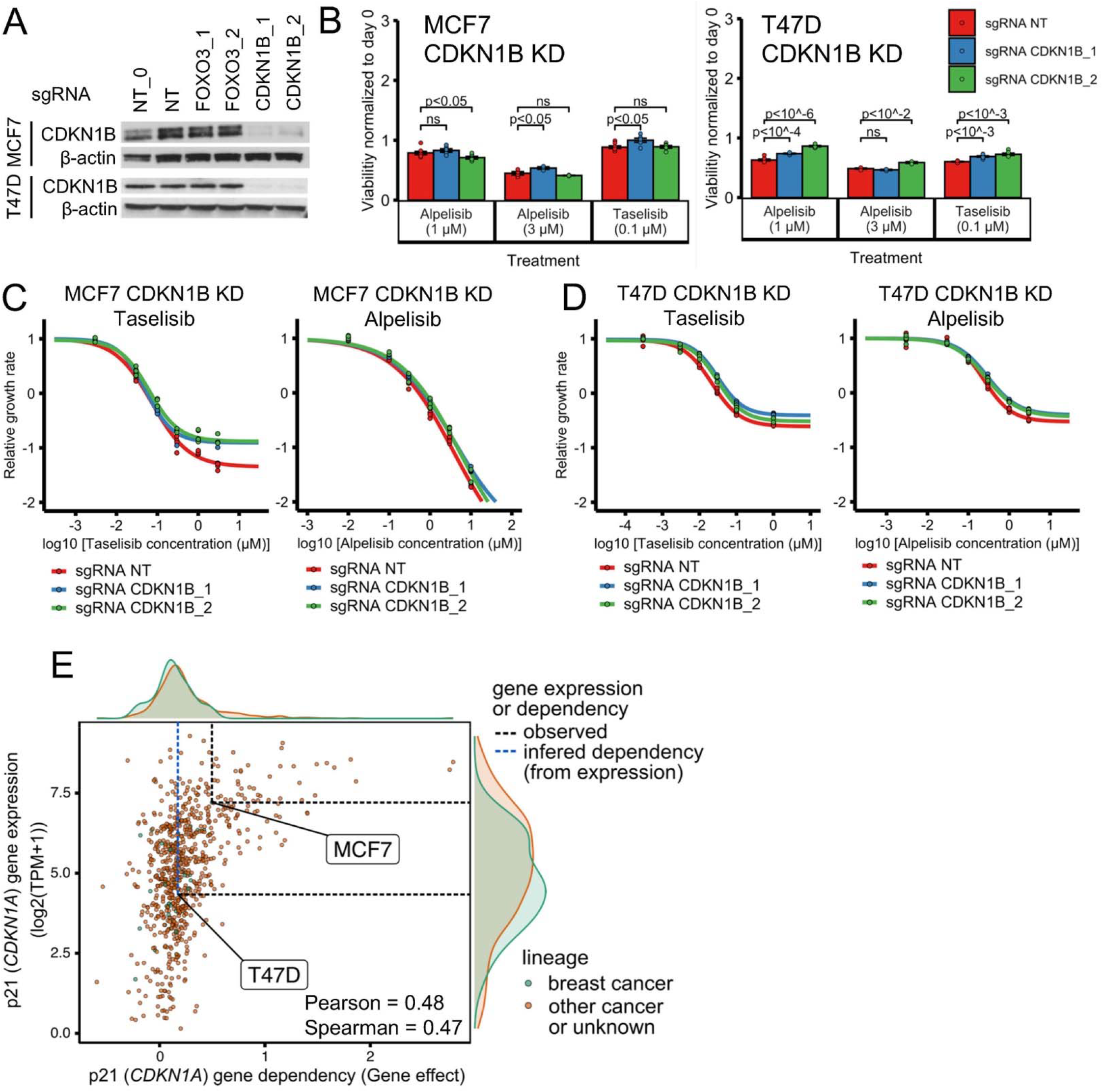
CDKN1B (p27) knockdown does not have a strong effect on PI3Kα inhibitor sensitivity in ER+ breast cancer cell line models. (A) Stable cell lines were generated using CDKN1B CRISPR knockdown and showed reduced proteins levels of CDKN1B. NT and NT_0 are two distinct non-targeting control guides. CDKN1B_1 and CDKN1B_2 are two distinct guides targeting CDKN1B. NT, NT_0, FOXO3_1, and FOXO3_2 are the same guides as in Fig. 4. (B-D) CDKN1B knockdown does not strongly reduce sensitivity to PI3Kα inhibitors alpelisib and taselisib in ER+ breast cancer cell lines MCF7 and T47D. The same experimental procedures as in panels B-D of Fig. 4 were used for panels B-D of this figure. Cell viability was measured before and after 6 days of drug treatment using CellTiter-Glo. Group comparisons between groups were done using an unpaired, two-sided t-test (panel B). Panels C and D calculate the relative growth rate with respect to DMSO control. (E) Sensitivity to p21 (CDKN1A) knockout in CRISPR/Cas-9 loss-of-function screens (DepMap) is strongly correlated with p21 gene expression. We hypothesize that the activity of p21 is behind the lack of a strong effect of p27 knockout on PI3Kα inhibitor sensitivity. Based on T47D’s p21 gene expression, the expected growth effect of p21 gene knockout on T47D is inferred to be smaller than that on MCF7. Sensitivity to gene knockout is measured using gene effect, which is such that gene effect=0 means no effect and gene effect=-1 is the median effect of known common essential genes. Inferred gene effect is shown with a dashed blue line; gene expression or gene effect from the dataset is shown with a dashed black line.

The lack of a strong effect of CDKN1B (p27) knockdown on PI3Kα inhibitor sensitivity can still be consistent with the model, if we hypothesize that only joint KD of CDKN1A (p21) and CDKN2B (p27) can accomplish what we observe in the model for the inactivation of the node *p21/p27*. To test the plausibility of this hypothesis, we looked at the gene expression and gene knockout effect of p21 and p27 using the Cancer Cell Line Encyclopedia (CCLE) and the CRISPR/Cas9 loss-of-function genetic screens of the Cancer Dependency Map (DepMap). Across all cell lines, p21 tended to show a tumor suppressor effect (*median(gene effect) = 0*.*20* for all 739 cell lines) and this effect was correlated with the cell line’s p21 (*CDKN1A*) gene expression (*Pearson correlation* = 0.48, *p* = 1.5 x 10^−43^; *Spearman correlation* = 0.47, *p* = 2.4 x 10^−41^). This was not the case for p27, which did not show a consistent tumor suppressor effect (*median(gene effect) = −0*.*01* for the for all 739 cell lines) and its effect was not correlated with its gene expression (*Pearson correlation* = 0.05, *p* = 1.9 x 10^−1^; *Spearman correlation* = 0.08, *p* = 2.7 x 10^−2^).

We also used the DepMap and CCLE database to look at whether we expect p21 knockout to show a difference in gene dependency between the MCF7 and T47D cell lines. MCF7 was in the high end of p21 gene expression (Fig. 5E; *log*_*2*_ *(TPM + 1) = 7*.*21*, robust z-score *z = 0*.*86*) and grew faster in response to CDKN1A gene knockout (Fig. 5E; *gene effect = 0*.*50*). T47D was in the low end of p21 expression (Fig. 5E; *log*_*2*_*(TPM + 1) = 4*.*33*, robust z-score *z = −0*.*54*) and, although it is not in the DepMap database, the inferred effect of p21 gene knockout on T47D was less than half of that for MCF7 (Fig. 5E; *median(gene effect) = 0*.*17* for the 23 cell lines within *±2*.*5%* of the *log*_*2*_ *(TPM + 1)* of T47D’s *CDKN1A* gene expression).

Overall, our experimental results suggest that CDK1NB (p27) knockdown does not have a strong effect on PI3Kα inhibitor sensitivity, in contrast with what the network model predicts. Based on computational analysis of gene expression and loss-of-function cell line databases, we hypothesize that the lack of a strong effect seen in CDK1NB knockdown could be explained by the activity of CDKN1A (p21); we expect MCF7 to show a stronger tumor suppressor effect than T47D in response to CDKN1A knockdown.

### RB1 knockdown reduces sensitivity to PI3Kα inhibitors in a cell line-specific manner

In the network model, PI3Kα inhibitors result in reduced cell proliferation by two mechanisms: by a decrease in E2F1 activity mediated by Rb or by a decrease in translational machinery mediated by mTORC1. As a result, the model predicts that RB1 knockdown would increase E2F1 activity and reduce the sensitivity to PI3Kα inhibition (Gómez Tejeda Zañudo, Scaltriti, and Albert 2017). To test this, we used T47D and MCF7 cell lines with sgRNA CRISPR KD of RB1 we previously generated (Wander et al. 2020).

Fig. 6A-B shows the drug response curve of RB1 KD T47D and MCF7 cell lines in response to PI3Kα inhibitor alpelisib. T47D RB1 KD cells showed a reduced sensitivity to alpelisib compared to control cells (Fig. 6A) and this reduced sensitivity was not seen in MCF7 cells (Fig. 6B). To further verify the reduced sensitivity to PI3Kα inhibitors in T47D RB1 KD cells, we used a long-term colony formation assay (Fig. 6C-D). This assay also indicates that T47D RB1 KD but not MCF7 RB1 KD cells (Fig. 6C) showed reduced sensitivity to alpelisib (Fig. 6D). In summary, we found that RB1 KD can reduce the sensitivity to PI3Kα inhibitor alpelisib, as predicted by the model, and that this effect was cell line-specific (observed in T47D but not MCF7).

**Fig 6.**
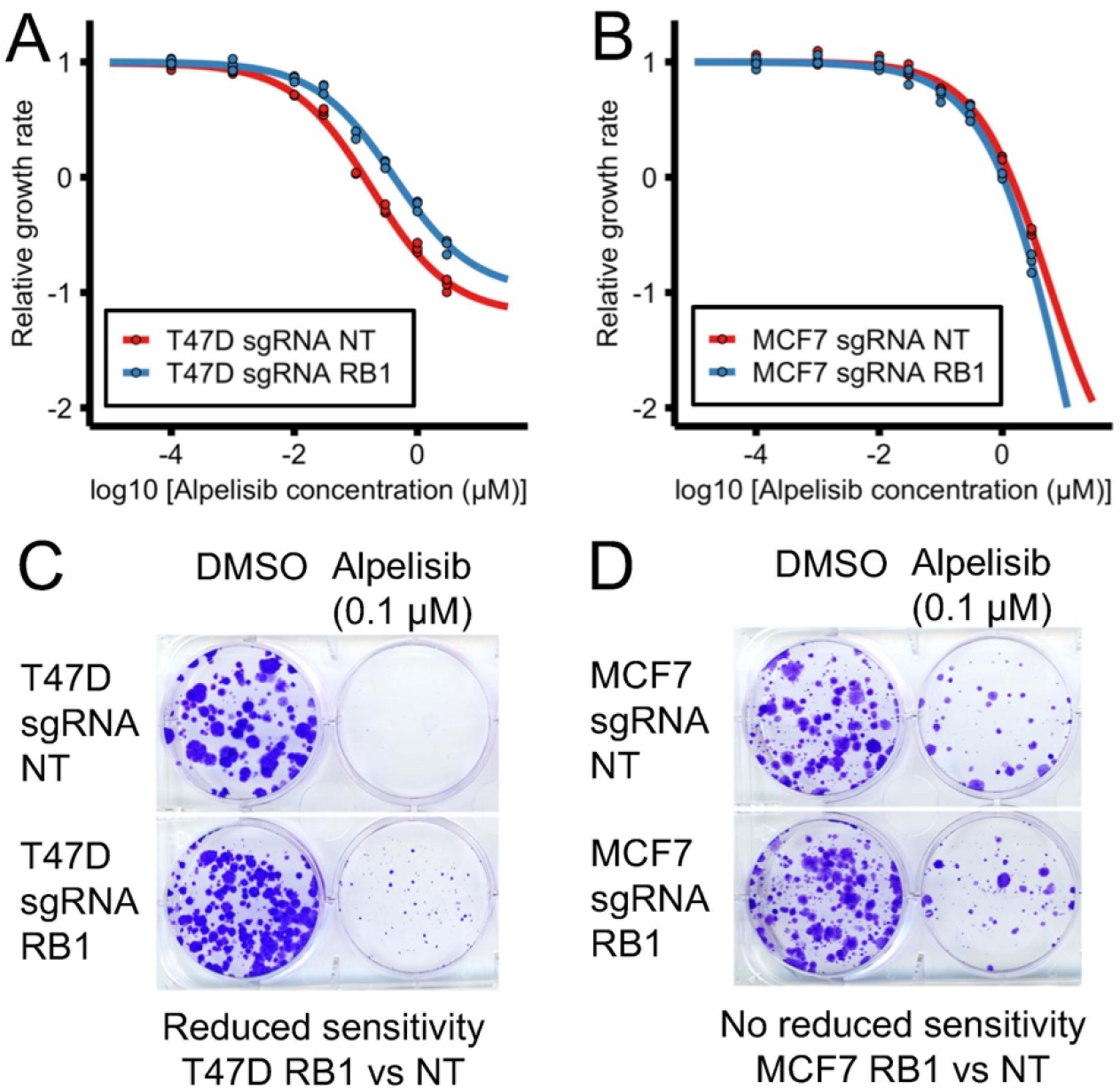
RB1 knockdown can reduce sensitivity to PI3Kα inhibitors in ER+ breast cancer cell line models. RB1 knockdown in T47D (panels A and C) but not MCF7 cells (panels B and D) results in a reduced sensitivity to PI3Kα inhibitor alpelisib in short-term and long-term assays. Cell viability was measured using a CellTiter-Glo assay before and after 6 days of drug treatment. Relative growth rate was calculated compared to the DMSO control treatment (panels A and B). Colony formation assay was performed after 30-40 days of drug treatment.

### Cell line-specific network models reproduce the efficacy of PI3Kα inhibitor drug combinations and resistance mechanisms

Our experimental results are in agreement with the model-predicted efficacy of combining PI3Kα inhibitors with BH3 mimetics and the reduced PI3Kα inhibitor sensitivity that results from FOXO3 knockdown or RB1 knockdown. However, our experimental results also exhibit cell line-specific aspects, which the model does not fully recapitulate. For example, MCF7 cells were sensitive to combined alpelisib and s63845 (Fig. 2) while T47D cells were only sensitive to combined alpelisib, s63845, and navitoclax (Fig. 3). Because of this, we generated cell line-specific network models for MCF7 and T47D.

The cell line-specific models are based on an updated version of the network model that incorporates new knowledge generated on ER+ breast cancer drug resistance since the previous model was published (e.g. (Hopkins et al. 2018; Donnella et al. 2018; Wander et al. 2020)) and the discrepancies from the model found by our experimental results (see Supplemental Text). The updated network model separates protein pairs (p21 and p27, BCL-2 and BCL-XL) that were previously denoted by a single combined node (p21/p27 and BCL2). The updated model includes a new node that denotes the transcriptional status of MYC targets (following the results from (Ilic et al. 2011; Donnella et al. 2018)) and assumes that the mTORC1 and Rb-E2F pathways do not contribute equally to the response to alpelisib (following our findings on RB1 and p27 knockdown). A condensed representation of the updated model is shown in Fig. 7A while a more complete representation of the model and the changes introduced is shown in Supp. Fig. 5A. This model describes the biological outcomes of proliferation and apoptosis using the multi-state nodes *Proliferation* and *Apoptosis* (each of which has 3 states) and their normalized propensities denoted by *Proliferation*_*norm*_ and *Apoptosis*_*norm*_ (which take values between 0 and 1).

**Fig 7.**
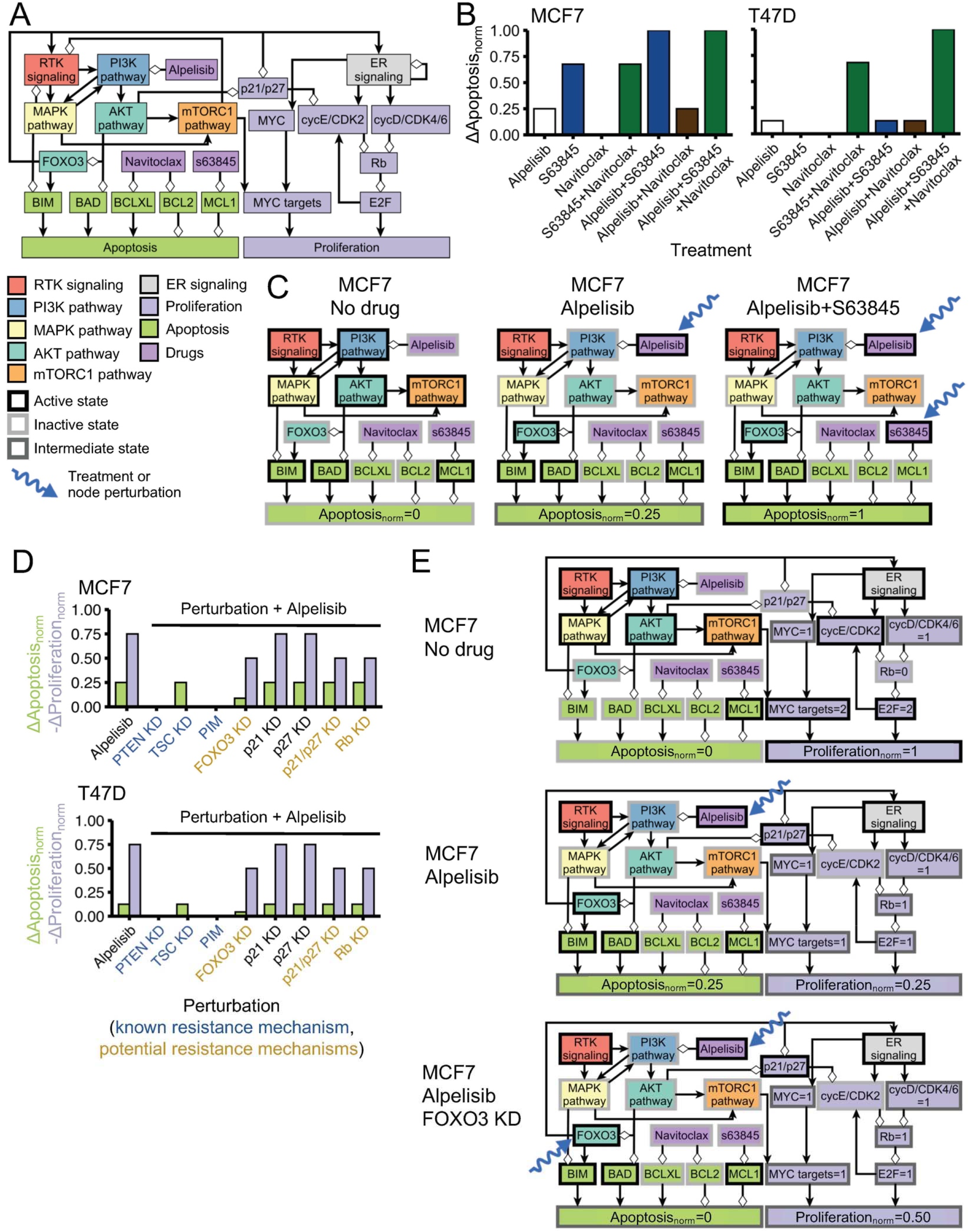
Cell line-specific network models reproduce the efficacy of combined PI3Kα inhibitors and BH3 mimetics and the effect of PI3Kα inhibitor resistance mechanisms in MCF7 and T47D. (A) Condensed representation of the network model in which the cell line-specific models are based. The condensed representation includes nodes denoting pathways (RTK signaling, PI3K pathway, MAPK pathway, AKT pathway, mTORC1 pathway, and ER signaling), biological outcomes (apoptosis, proliferation), and drugs or proteins of interest for this work (e.g. alpelisib, s63845, navitoclax, FOXO3, p21/p27, and Rb). A more complete representation of the network model is shown in Supp. Fig. S4. (B-C) MCF7-specific and T47D-specific network models reproduce the difference in cell death responses to BH3 mimetics, alpelisib, and their combination seen in these cell lines. The model results in panel B are comparable to the experimental results in Fig. 2E (MCF7) and 2C (T47D) (see Supp. Fig S6). We show the state of the nodes that influence Apoptosis in response to alpelisib, BH3 mimetics, and their combination for the case of MCF7 (panel C) and T47D (Supp. Fig S5B). (D-E) Cell line-specific network models reproduce the drug resistance effect of PI3Kα inhibitor resistance mechanisms. Simulations in which a PI3Kα inhibitor resistance mechanism is active show an increased survivability (reduced *Proliferation* and/or increased *Apoptosis*) compared to the case of alpelisib alone (panel D). We show the state of the nodes that influence *Apoptosis* and *Proliferation* in response to alpelisib, FOXO3 knockdown, and their combination for the case of MCF7 (panel E) and T47D (Supp. Fig S5D). The models encode the biological outcomes of cell death and proliferation using *Apoptosis* _*norm*_ and *Proliferation* _*norm*_, respectively, which are a weighted and normalized (between 0 and 1) measure of the state of the nodes *Apoptosis* and *Proliferation*. Starting from a cancerous state of each model, we perform 10,000 simulations in which the specified treatment or perturbation is maintained throughout the simulation, and obtain the average value of *Apoptosis* _*norm*_ and Proliferation _*norm*_ at the end of the simulations. *Δ Apoptosis* _*norm*_ and *Δ Proliferation* _*norm*_ of a perturbation denote the difference with respect to the case of no perturbation (*Apoptosis* _*norm*_ = 0, *Proliferation* _*n orm*_ = 1), and are such that decreased survivability means an increase in *Δ Apoptosis* _*norm*_ or -*Δ Proliferation* _*norm*_.

The MCF7-specific and T47D-specific network models differ only in three aspects: (1) the node denoting basal expression of BCL-XL is active in T47D but not MCF7 (in agreement with Fig. 3E), (2) the node denoting basal expression of p21 is more active in MCF7 than in T47D (in agreement with Fig. 5E), and (3) the contribution of the lowest active *Apoptosis* state to *Apoptosis*_*norm*_ is larger in MCF7 than T47D (i.e., *Apoptosis* = 1 results in *Apoptosis*_*norm*_ = 0.25 in MCF7 and *Apoptosis*_*norm*_ = 0.125 in T47D) (in agreement with Figs. 2D and 3D).

The cell line-specific models’ cancerous states (which are steady states) and the time course of their response to alpelisib share many similarities with that of the previous model. The cancerous states have active RTK, PI3K, MAPK, AKT, mTORC1, and ER signaling, which result in high survivability (*Proliferation*_*norm*_ *=1* and *Apoptosis*_*norm*_ *= 0*). These states can be primed for cell death, a common feature of cancer cells (Ni Chonghaile et al. 2011; Montero et al. 2015), by having pro-apoptotic protein BIM active but an anti-apoptotic protein counteracting its effect (BCL-2, BCL-XL, or MCL1 in the updated model). The time course of the MCF7-specific model in response to alpelisib starting from a cancerous state is shown in Supp. Fig S4B. In response to alpelisib, there is a quick inactivation of the MAPK, AKT, and mTORC1 pathways in both cell line-specific models, followed by the activation of FOXO3, which transcriptionally upregulates HER3 and ESR1. The inhibition of the mTORC1 pathway downregulates the node *MYC targets*, which results in reduced proliferation (*Proliferation*_*norm*_ *= 0*.*25*) in both models. The activation FOXO3 and inactivation of AKT results in the activation of pro-apoptotic proteins BIM and BAD, which result in a small increase in cell death (*Apoptosis*_*norm*_ *= 0*.*25* in MCF7 and *Apoptosis*_*norm*_ *= 0*.*125* in T47D).

These cell line-specific network models reproduce the observed difference in cell death response to BH3 mimetics and their combination with alpelisib in each cell (see Fig. 7B and Supp. Fig S6, where *ΔApoptosis*_*norm*_ is the difference with respect to the case of no perturbation). In the MCF7 model, alpelisib alone has a small effect on apoptosis (*ΔApoptosis*_*norm*_ *= 0*.*25*), s63845 alone has larger effect on apoptosis (*ΔApoptosis*_*norm*_ *= 0*.*68*), and combined alpelisib and s63845 has the largest effect (*ΔApoptosis*_*norm*_ *= 1*) (Fig. 7B-C). In the T47D model, alpelisib alone has a small effect on apoptosis (*ΔApoptosis*_*norm*_ *= 0*.*125*), s63845 or navitoclax alone have no effect (*ΔApoptosis*_*norm*_ *= 0*), their combination has a larger effect (*ΔApoptosis*_*norm*_ *= 0*.*68*), and their combination with alpelisib has the largest effect (*ΔApoptosis*_*norm*_ *= 1*) (Fig. 7B and Supp. Fig S5B).

The cell line-specific models recapitulate how FOXO3 knockdown reduces sensitivity to alpelisib in both MCF7 (Fig. 7D-E) and T47D (Fig. 7D and Supp. Fig S5D), and the stronger reduction in apoptosis seen in MCF7 (Fig. 7D). For the case of Rb knockdown and p21/p27 knockdown, the models reproduce the observations that their effect on sensitivity to alpelisib is small (*Proliferation*_*norm*_ *= 0*.*25* to *Proliferation*_*norm*_ *= 0*.*5*) and that p27 knockdown alone does not have a strong effect (no change in *Proliferation*_*norm*_), and also incorporate the hypothesis that the effect of p21 and p27 requires knockdown of both p21 and p27 (Fig. 7D).

The MCF7-specific and T47D-specific network models recapitulate the PI3K inhibitor resistance mechanisms and drug combinations that the previous model also did. For example, knockdown of PTEN, knockdown of TSC, or high expression of PIM make the cell insensitive to alpelisib (*Proliferation*_*norm*_ *= 0*.*25* with alpelisib changes to *Proliferation*_*norm*_ *=1* with the addition of any of these resistance mechanisms), and the addition of fulvestrant to alpelisib results in a larger effect than any of them alone (*Proliferation*_*norm*_ *= 0*.*25* with alpelisib or fulvestrant alone changes to *Proliferation*_*norm*_ *=0* with both) (Fig. 7D and Supp. Fig S5A and C). Additional resistance mechanisms to alpelisib or PI3K inhibitors that both models reproduce include high expression of SGK1/PDK1 (Castel et al. 2016), the HER3 microenvironment (Kodack et al. 2017), high activity of IGF1R and insulin (Hopkins et al. 2018), upregulation of ESR1 expression (Bosch et al. 2015), and high MYC activity (Ilic et al. 2011) (Supp. Fig S5C). The models also recapitulate the resistance mechanisms of other drugs. For example, Rb knockdown is a resistance mechanism to palbociclib (Herrera-Abreu et al. 2016) (*Proliferation*_*norm*_ = 0.25 with palbociclib changes to *Proliferation*_*norm*_ *=1* with added Rb knockdown) but has little effect in the presence of fulvestrant (Wander et al. 2020) (*Proliferation*_*norm*_ = 0.25 with fulvestrant to *Proliferation*_*norm*_ = 0.5 with the addition of Rb knockdown) (Supp. Fig S5C).

Overall, the cell line-specific network models we developed are able to reproduce the current knowledge of PI3Kα inhibitor resistance mechanisms and drug combinations, including the new results from this work on the efficacy of the combination of alpelisib with BH3 mimetics and the reduced sensitivity to PI3Kα inhibitors as a result of FOXO3 knockdown.

## Discussion

In this work, we experimentally validated several PI3Kα inhibitor drug resistance and sensitivity factors predicted by a network-based mathematical model of oncogenic signaling in ER+ *PIK3CA* mutant breast cancer (Fig. 1). We experimentally verified (i) the efficacy of combining PI3Kα inhibitor alpelisib with BH3 mimetics (MCL1 inhibitor s63845 alone or in combination with BCL-XL/BCL-2 inhibitor navitoclax, in a cell line-specific manner), a promising PI3Kα inhibitor drug combination (Figs. 2 and 3) and (ii) the reduced sensitivity to PI3Kα inhibitors (alpelisib and taselisib) caused by FOXO3 knockdown, a novel candidate resistance mechanism for PI3Kα inhibitors (Fig. 4). We additionally found discrepancies with the model, in particular, our experiments found that the effect of p27 knockdown and RB1 knockdown did not result in the expected reduced sensitivity or had only a small cell line-specific effect (Figs. 5 and 6). Based on these experiments, we developed updated and cell line-specific network models that can recapitulate our new results and other recent work on PI3Kα inhibitor drug resistance.

Our combined mathematical modeling and experimental approach follows the model/experiment refinement cycle that is a cornerstone of systems biology research (Coutant et al. 2019; Alberghina and Westerhoff 2007), and applies it to the field of cancer drug resistance. In this approach, cancer biology and clinical knowledge are integrated to form a model, experiments are performed to test and refine the model, new knowledge is subsequently generated, and the cycle then repeats itself. This completion of a model/experiment cycle is a step that is necessary for the development of predictive and validated mathematical models in cancer and systems biology, yet it is rarely accomplished (Coutant et al. 2019; Brady and Enderling 2019; Alberghina and Westerhoff 2007).

An important result from this work is that the combination of PI3Kα inhibitor alpelisib with BH3 mimetics is synergistically efficacious and that this combination leads to an increase in apoptosis compared to each agent alone (Figs. 2 and 3). We find that the ideal choice of BH3 mimetics in the combination strategy depends on which anti-apoptotic BCL-2 family members (e.g. MCL1, BCL-XL, and BCL-2) are highly expressed in a given cancer cell. For the case of T47D cells, which have high BCL-XL expression, alpelisib was synergistically efficacious only when combined with MCL1 inhibitor s63845 and BCL-XL/BCL-2 inhibitor navitoclax. Meanwhile, for MCF7 cells, which have low BCL-XL expression, alpelisib was synergistically efficacious when combined with s63845 alone.

These cell line-specific results highlight an important lesson for the use of alpelisib and BH3 mimetics in the clinical setting of breast cancer: fully taking advantage of this drug combination will require determining the relevant anti-apoptotic BCL-2 family dependence in the tumor. This could be accomplished by directly quantifying these BCL-2 family members in the tumor and/or by using functional genomic approaches such as dynamic BH3 profiling (Montero et al. 2015; Montero et al. 2019; Bhola et al. 2020). In line with this view, recent clinical trials on breast cancer tumors with high BCL-2 expression have shown promising results with the combination of approved breast cancer therapies with the BCL-2 inhibitor venetoclax (Whittle et al. 2020; Lok et al. 2019). Our results suggest a similar approach using the combination of alpelisib and tumor-specific BH3 mimetics determined through dynamic BH3 profiling.

Another key result from this work is that FOXO3 knockdown reduces the sensitivity to PI3Kα inhibitors, thus revealing a novel candidate resistance mechanism (Fig. 4). FOXO3 exemplifies the unique insights that can be obtained from network models and their ability to integrate the context-dependent role of cancer-related genes. FOXO3 is known for its dual role as tumor suppressor and oncogene, acting as one or the other depending on the context. Previous work had highlighted the role of FOXO3 as an oncogene through its PI3K-inhibitor-induced feedback activation and the resulting upregulation of ESR1 and HER3. The upregulation of these receptors leads to increased survival in microenvironments that favor their signaling (Kodack et al. 2017; Bosch et al. 2015). Our network modeling and experimental findings show that the tumor suppressor effect of FOXO3 can dominate.

Overall, we applied a systems biology approach combining a network-based mathematical model and its experimental validation to tackle the problem of PI3Kα inhibitor drug resistance in ER+ *PIK3CA* mutant breast cancer (Fig. 1). The result was the experimental validation of BH3 mimetics as an efficacious drug combination and of FOXO3 knockdown as a drug sensitivity factor and potential resistance mechanism. These results led to a refinement of the model and the development of cell line-specific network models that recapitulate the observed cell line-specific effects and the current knowledge on drug resistance to PI3Kα inhibitors. The validated drug combinations and sensitivity factors of our approach illustrate how the use of network modeling and systems biology approaches can contribute to overcoming the challenge of cancer drug resistance.

## Acknowledgements

J.G.T.Z. would like to thank all members of the Wagle lab and all members of the Stand Up To Cancer – National Science Foundation Drug Combination Convergence Research Team for many useful discussions. J.G.T.Z. would also like to thank Stand Up To Cancer and V foundation staff, grant reviewers, and scientific leadership for organizing and making possible multiple meetings, which shaped and enriched this work. J.G.T.Z. acknowledges Christian Meyer and Vito Quaranta for their help in quantifying drug combination synergy using MuSyC. J.M. acknowledges support from the Ramon y Cajal Programme, Ministerio de Economia y Competitividad (RYC-2015-18357). J.M. and C.A. acknowledge the CELLEX foundation. This work was supported by the National Science Foundation grants PHY 1545832 (to R.A.), PHY 1545839 (to A.G.L. and N.W.) and 1545853 (to M.S. and G.X.), the Stand Up to Cancer Foundation, and Stand Up to Cancer Foundation/The V Foundation Convergence Scholar Awards to J.G.T.Z. (D2015-039), J.M. (D2015-037), and P.M..

## Methods

### Cell culture

293T, T47D and MCF7 cells were purchased from American Type Culture Collection (ATCC). 293T cells were cultured in DMEM (GIBCO #11995-065) with 10% fetal bovine serum (Gemini #100-119). T47D cells were cultured in Phenol Red-free RPMI-1640 (GIBCO #11835-030) with 10% FBS. MCF7 cells were cultured in Phenol Red-free MEM-*α* (GIBCO #41061-029) with 10% FBS.

### Drug treatment

Drugs that were used to treat cells include taselisib (Selleck Chemicals # S7103), alpelisib (Selleck Chemicals # S2814), s63845, everolimus (Selleck Chemicals # S1120), navitoclax (Selleck Chemicals # S1001), fulvestrant (Sigma Aldrich # I4409), ipatasertib (Thermo Fisher Scientific # NC0742252), palbociclib (Selleck Chemicals #S1116).

### Generation of plasmids and engineered cells

We cloned sgRNA into lentiCRISPRv2 vector (gift from Dr. Adam Bass’s laboratory) and lentivirus was produced in 293T cells using psPAX2 and pCMV-VSV-G plasmids. T47D or MCF7 cells were infected with lentivirus to derive stable cell lines expressing sgRNA to target FOXO3 (FOXO3_1: CGCCGACTCCATGATCCCCG, FOXO3_2: GGAAGAGCGGAAAAGCCCCC) or CDKN1B (CDKN1B_1: GGAGAAGCACTGCAGAGACA, CDKN1B_2: GGACCACGAAGAGTTAACCC) or non-targeting (NT) sgRNA controls (NT_0: GTATTACTGATATTGGTGGG, NT: ACGGAGGCTAAGCGTCGCAA). The sequences for the guides for FOXO3 and CDKN1B were obtained from the DepMap 18Q3 release. Cells were selected by puromycin (Life Technologies #A1113803). RB1 and CRISPR non-targeting guide cells used as control for RB1-related experiments were obtained as a gift from Flora Luo and the Garraway laboratory, and have been previously characterized (Wander et al. 2020).

### Kill curves and CellTiter-Glo viability assay

Cells were plated in 96-well tissue culture ViewPlate (Perkin Elmer # 6005181) on Day 1 and treated with drug on Day 2. Media with or without drugs was refreshed on Day 5. On Day 8, cells were equilibrated to room temperature, media was removed, and cells were lysed in a mixture of 50 µL media and 50 µL CellTiter-Glo 2.0 reagent (Promega # G9243) per well. Plates were then incubated on an orbital shaker for 2 mins. Following another 10 mins of incubation at room temperature to stabilize signal, luminescence was recorded to measure cell viability on Infinite M200 Pro microplate reader (Tecan).

### Western blotting

Cell lysates were prepared in RIPA buffer (Sigma Aldrich # R0278) supplemented with dithiothreitol (Life Technologies # 15508013), phenylmethane sulfonyl fluoride (Sigma Aldrich # P7626), protease inhibitor cocktail (Sigma Aldrich # P8340) and phosphatase inhibitor tablets (Roche # 4906845001). NuPAGE 4-12% Bis-Tris Midi protein gels (Invitrogen # WG1402A) were used for western blot. After electrophoresis, gel was transferred using Trans-Blot Turbo PVDF or nitrocellulose transfer pack (Bio-Rad # 1704158 and # 1704156). Membranes were probed with specific primary antibodies at 4 °C overnight and then incubated with corresponding HRP-conjugated secondary antibodies. We use Pierce ECL Plus Western Blotting Substrate (Thermo Fisher Scientific # 32132×3) as developing reagent. Primary antibodies used in this study are: FOXO3 (CST #2497S), CDKN1B (CST #3698S), p-AKT (CST #12694S), pS6 (CST #4857S), MCL (CST #5453S), PARP (CST #9532S), cleaved PARP (CST #5625S), Actin (C4) (SC-47778). Secondary antibodies are from Invitrogen (anti-mouse A16090, anti-rabbit 32260, anti-goat 81-1620).

### Colony formation assay

1,000-10,000 cells were plated in 6-well plates on Day 1 and treated on Day 2. Media was refreshed every 3-4 days until crystal violet staining. On the day of staining, cells were fixed with ice-cold 100% methanol for 10 minutes and then incubated with 0.5% crystal violet solution (Sigma Aldrich #C6158) in 25% methanol at room temperature for 10 minutes.

### Cell death assays

Cells were treated as indicated and stained with fluorescent conjugates of annexin-V and PI (1 µg/ml final concentration) and analyzed on a FACSCanto machine (BD, Franklin Lakes, NJ, USA). Annexin V was prepared as previously described (Brumatti et al., Methods 2008). Viable cells are Annexin-V negative and PI negative, and cell death is expressed as 100% – viable cells.

### Dynamic BH3 profiling

Dynamic BH3 profiling was performed as previously described in detail (Montero et al. Cancer Discovery 2017). ER+ breast cancer cell lines were incubated in RPMI with 10% FBS for T47D and DMEM with 10% FBS for MCF7, at different times and drug concentration as indicated. To perform DBP in cell lines, 2 × 10^4^ cells/well were used. We used BIM BH3 peptide concentrations of 0.01, 0.03, 0.1, 0.3, 1, 3, and 10 μM. We used a FACS-based BH3 profiling to perform the analysis, cytochrome c release was measured after a 60 minutes incubation of digitonin-permeabilized cells with BH3 peptides, as previously described (Ryan et al., Biol. Chem 2016). % priming stands for the % cytochrome c released for each BH3 concentration tested. *Δ*% priming stands for the maximum difference in % priming between treated (drug) and non-treated (no drug) cells among the BH3 concentrations tested.

### Synergy scores

We quantify drug combination synergy using MuSyC (Meyer et al. 2019). MuSyC calculates the synergy score from drug combination kill curves and considers separately the effect of efficacy and potency. In this work, we focus on the synergy score of efficacy (*β*_*obs*_). *β*_*obs*_ measures the percentage change in drug response of the drug combination when compared to that of each drug alone (at the highest concentration used for each drug alone and their combination). The synergy score of efficacy *β*_*obs*_ is such that *β*_*obs*_ *>0* means the drugs are synergistic, *β*_*obs*_ *<0* that they are antagonistic, and *β*_*obs*_ *=0* means the drugs show no synergistic or antagonistic effect.

### Analysis of loss-of-function genetic screens, gene expression, and protein expression in cell line datasets

Analysis of the loss-of-function genetic screens from the Cancer Dependency Data (DepMap) and expression data from the Cancer Cell Line Encyclopedia (CCLE) followed a similar approach as in (Painter et al. 2020). Gene dependency data from CRISPR knockout (Avana data set), RNAi knockdown (Achilles and DRIVE data sets), gene expression (Ghandi et al. 2019) (RNA-seq from the CCLE), relative protein expression (Nusinow et al. 2020) (quantitative mass-spectroscopy proteomics from CCLE), and associated cell line annotations were taken from the DepMap 20Q1 Public data release (Dempster et al. 2019; DepMap 2020) (https://depmap.org/portal/download). Dependency for each gene and cell line is obtained using the CERES algorithm (Meyers et al. 2017) (for CRISPR knockout) or the DEMETER2 algorithm (McFarland et al. 2018) (for RNAi knockdown). Gene dependency is measured using the gene effect, which measures the impact of CRISPR knockout or RNAi knockdown of a gene on cell viability. The gene effect is normalized for each cell line so that a gene effect of 0 is the median dependency score of a negative control gene and a gene effect of - 1 is the median dependency score of pan-essential genes (Dempster et al. 2019).

To infer the gene effect of MCL1 and p21 (*CDKN1A*) in T47D (a cell line not included in the DepMap CRISPR knockout screens), we use the gene effect of cell lines in which the gene of interest has a similar expression as in T47D. The inferred gene effect is defined as the median gene effect of cell lines within *±2*.*5%*of the *log*^*2*^ *(TPM + 1)* of T47D’s expression of the gene of interest. In particular, we use *CDKN1A* expression to infer p21 gene effect and *BCL2L1* expression to infer MCL1 gene effect. This inference is based on the observation that the gene effect of MCL1 and the gene expression of *BCL2L1* are correlated across cell lines (*Pearson correlation* = 0.45, *p* = 1.9 x 10^−37^; *Spearman correlation* = 0.47, *p* = 7.2 x 10^−41^), and similarly for the gene effect of p21 and the gene expression of *CDKN1A* (*Pearson correlation* = 0.48, *p* = 1.5 x 10^−43^; *Spearman correlation* = 0.47, *p* = 2.4 x 10^−41^). For the case of MCL1, the inferred MCL1 gene effect is *-0*.*40* (*median(gene effect)* = −0.40 for the 57 cell lines within ±2.5% of the *log*_*2*_ *(TPM + 1)* of T47D’s *BCL2L1* gene expression). For the case of p21, the inferred p21 gene effect is 0.17 (*median(gene effect)* = 0.17 for the 23 cell lines within ±2.5% of the *log*_*2*_ *(TPM + 1)* of T47D’s *CDKN1A* gene expression).

### Statistical analysis

All statistical analysis was performed using R (version 3.6.0). Pearson and Spearman correlations and *p* values were calculated using the *cor*.*test* function from the *stats* package using a two-sided test. Statistical comparisons in cell viability were performed using a two-sample, two-sided unpaired t-test using the *t*.*test* function from the *stats* package. A *p* value of less than 0.05 was considered to be statistically significant. Robust z-scores *z* for a value *x*_*j*_ in a distribution *D = [x*_1_, …, *x*_*n*_] were obtained by subtracting the median (*median*) and dividing by the median absolute deviation (*mad*), that is, *z(x*_*j*_ *) = [x*_*j*_ *-median(D)]/mad(D)*.

### Dynamic network modeling

Our dynamic network modeling approach follows the methodology in (Gómez Tejeda Zañudo, Scaltriti, and Albert 2017), which uses a type of dynamic network model known as discrete dynamic network model. In a discrete dynamic network model, each node *i* in the directed network that underlies the model has an associated variable *σ*_*i*_ that can only take a discrete number of states. In the simplest case, the Boolean case, a node variable can only take the two states: *1* (active) or 0 (inactive). The state of a variable *σ*_*i*_ is determined using a regulatory function *f*_*-i*_, which depends on the variable of a node *j* only if there is a directed interaction from *j* to *i* (*j → i*).

The state of the system *∑(t) = [σ*_1_*(t), σ*_2_*(t), …, σ* ^*N*^*(t)]*, where *σ*_*i*_ *(t)* is the state of variable *σ*_*i*_ at time *t*, changes in time following a general asynchronous updating scheme (Garg et al. 2008; Saadatpour, Albert, and Albert 2010). In general asynchronous updating, the state of the system is updated in discrete time units by (1) choosing one randomly selected node *k* at each time step *t* (with update probability *p*_*k*_) and updating its state based on the state of the system at the previous time step (*σ*_*k*_*(t) = f*_*k*_*(∑(t - 1)*) and (2) transferring the node state from the previous time step of the rest of the nodes (*σ* _*l*_*(t) =σ*_l_*(t - 1),l* ≠*k*).

### Model simulationssss

Model simulations were performed using similar methodology as in (Gómez Tejeda Zañudo, Scaltriti, and Albert 2017). The simulations of the discrete network models were done by mapping the discrete network model into a Boolean model, which was then simulated using the BooleanDynamicModeling Java library, which is available on GitHub (https://github.com/jgtz/BooleanDynamicModeling). To simulate multi-level nodes, we use a Boolean variable to denote each level greater than 1. For example, for a 3-level node with states 0, 1, and 2, we have 2 variables (Node and Node_2), and for a 4-level node we have 3 variables (Node, Node_2, and Node_3). The updating probabilities are chosen by categorizing nodes into either a fast or slow node, according to whether activation of the node denotes a (fast) signaling event or a (slow) transcriptional or translational event. The regulatory functions of all the nodes are indicated and explained in Supplemental Text and Supplemental File 1. We perform 10,000 simulations in each modeled scenario. The number of time units is 25 for all simulations, where a time unit is equal to the average number of time steps needed to update a slow node. More details on the network model and the model simulations can be found in the Supplemental Text and Supplemental File 1.

The code used to simulate the model is available on GitHub (https://github.com/jgtz/BreastCancerModelv2), and includes a Jupyter notebook that generates all the model results. To allow easy reuse of the model, we use bioLQM (Naldi 2018) (version 0.6.1, https://github.com/colomoto/bioLQM) to make the model available in Systems Biology Markup Language (SBML) format on the GitHub repository.

## List of Supplemental Material

**Supplemental File 1**

**Supplemental Text**

**Supplemental Figures S1-S6**

