## Supplemental Text for "Cell line-specific network models of ER+ breast cancer identify PI3Kα inhibitor sensitivity factors and drug combinations"

### **Updated network model**

The updated network model used for the cell line-specific network models of MCF7 and T47D is based on our previous model (Gómez Tejeda Zañudo, Scaltriti, and Albert 2017). The main modifications to the network model are (i) the addition of nodes to explicitly represent gene products that were previously a single combined node ($p21\_T$ and $p27\_T$ instead of $p21/p27\_T$, $BCL2$and $BCLXL$ instead of $BCL2$) and the addition of drugs that target some of these nodes ($S63845$, $Navitoclax$), (ii) the addition of nodes that denote microenvironments ($ER\_microenvironment$, $HER3\_microenvironment$) that were previously implicitly included, (iii) the addition of the node $MYC\_targets$, (iv) the addition of nodes to explicitly denote the transcriptional status of certain genes of interest ($BCLXL\_T$, $MCL1\_T$, $MYC\_T$, $Rb\_T$, $PTEN\_T$), (v) a negative interaction from $ER transcription$ to $ESR1$, and (vi) minor modifications to the network structure based on a re-evaluation of the evidence used for those interactions and our experimental results.

In addition to these changes in the network structure, the updated model modified the regulatory function of multiple nodes. (The regulatory function of a node determines its next state as a function of the state of its regulators. Because we denote each node level with a Boolean variable, the regulatory function of each Boolean variable can be expressed as a logical rule.) The great majority of these changes were minor and a direct consequence of the addition of new nodes and their edges. The most significant change was in the regulatory functions for the nodes $Apoptosis$, $Proliferation$, $pRb$, and $E2F$. Here we give an overview of the updated model and the most significant changes. More details on the modifications to each rule of the model can be found in Supplemental File 1.

#### Explanation of node names

ESR1 – the transcript of the *ESR1* gene, which encodes the estrogen receptor.

ER - estrogen receptor status of the cell.

ER_transcription - ESR1's activity in regulating the transcription of its target genes. Depends on the activity of ESR1 and its cofactors.

ER_transcriptional_feedback_1 - Encodes for the biological outcome of high activity of ER_transcription and the negative interaction from ER transcription to ESR1

ER_transcriptional_feedback_2 - Same as ER_transcriptional_feedback_1, but has memory of its past activation through a positive self-loop.

ER_microenvironment - Estrogen-rich cellular microenvironment. When combined with upregulation of ESR1 it can result in high activity of ER_transcription.

HER2- human epidermal growth factor receptor 2.

HER3 - human epidermal growth factor receptor 3.

HER3_T - transcript of HER3.

HER2/3 – heterodimer of HER2 and HER3.

HER3_microenvironment - Neuregulin-rich cellular microenvironment. Together with upregulation of HER3 it can result in high activity of HER2/3.

IGF1R – insulin-like growth factor 1 receptor (IGF1R) and insulin receptor (INSR).

IGF1R_T – transcript of IGF1R or INSR.

PIP3 – phosphatidylinositol – (3,4,5) –triphosphate.

PI3K – phosphatidylinositol – 4,5- biphosphate 3-kinase, encoded by the gene *PIK3CA.*

PTEN – phosphatase and tensin homolog.

PTEN_T – transcript of PTEN.

AKT – protein kinase B.

RAS – Ras family small GTPase.

MAPK – merged node that includes RAF- rapidly accelerated fibrosarcoma kinase family, MEK - mitogen-activated protein kinase kinase, and ERK- extracellular signal-regulated kinase.

FOXA1 – forkhead box protein A1.

PBX1 – pre-B cell leukemia transcription factor 1.

KMT2D – histone-lysine N methyltransferase 2D.

MYC - transcription factor encoded by the proto-oncogene *c-myc.*

MYC_T - transcript of MYC.

MYC_targets - Genes regulated by MYC’s transcriptional regulatory activity and mTORC1-pathway-dependent protein translation

PDK1 – 3-phosphoinositide dependent protein kinase 1, encoded by the gene *PDPK1.*

PDK1_pm - 3-phosphoinositide dependent protein kinase 1 localized in the plasma membrane.

SGK1 – serum and glucocorticoid-induced kinase 1.

SGK1_T – transcript of serum and glucocorticoid-induced kinase 1.

PIM – Pim-1 Proto-Oncogene, Serine/Threonine Kinase (PIM1), Pim-2 Proto-Oncogene, Serine/Threonine Kinase (PIM2), and Pim-3 Proto-Oncogene, Serine/Threonine Kinase (PIM3). Denotes both the transcript of these genes and their signaling activity.

CDK4/6 - cyclin dependent kinase 4/6.

cyclinD - cyclin D1.

cycD/CDK4/6 – complex formed by cyclin D1 and cyclin dependent kinase 4/6.

cycE/CDK2 – complex formed by cyclin E and cyclin dependent kinase 2.

cycE/CDK2_T – transcriptional status of cyclin E and cyclin dependent kinase 2.

pRb – Inactive state of retinoblastoma (Rb) family proteins, in which Rb proteins are unbound from E2F family members.

Rb_T - Transcript of retinoblastoma (Rb) family proteins.

E2F - transcriptional activator members of the E2F family.

p21 – WAF1/CIP1, cyclin-dependent kinase inhibitor 1, encoded by the gene *CDKN1A.*

p21_T - Transcript of p21.

p27- KIP1, cyclin-dependent kinase inhibitor 1B, encoded by the gene *CDKN1B.*

p27_T - Transcript of p27.

p21/p27 - Combined activity of p21 and p27.

p21/p27_T - Combined transcriptional status of p21 and p27.

mTORC1 - mechanistic target of Rapamycin complex 1.

mTORC2_pm - mechanistic target of Rapamycin complex 2 localized in the plasma membrane.

mTORC2 - mechanistic target of Rapamycin complex 2 not localized in the plasma membrane and possibly localized in the mitochondria.

TSC – tuberous sclerosis complex 1 (TSC1) and tuberous sclerosis complex 2 (TSC2).

PRAS40 - proline-rich Akt substrate of 40 kDa, a component of mTORC1.

FOXO3 - forkhead box O3 protein.

FOXO3_Ub – forkhead box O3 protein degraded through the ubiquitin proteasome pathway.

S6K – p70 ribosomal S6 kinase.

EIF4F - eukaryotic initiation factor 4A (EIF4A), eukaryotic initiation factor 4G (EIF4G), eukaryotic translation initiation factor 4E (EIF4E), and eukaryotic translation initiation factor 4E binding protein 1 (4EBP1).

Translation – processes related to ribosome translation, cap-dependent translation.

Proliferation – Propensity of the cell to commit to cell cycle progression. Focuses on two requirements: cell cycle transition from G1 to S (through E2F), and the transcriptional status of MYC targets.

BCL2- B-cell lymphoma 2 protein. Part of the anti-apoptotic BCL-2 family.

BCL2_T – B cell lymphoma 2 transcript

MCL1 – Encoded by the gene *MCL1*. Part of the anti-apoptotic BCL-2 family.

MCL1_T - MCL1 transcript

BCLXL – B-cell lymphoma-extra large, encoded by the gene *BCL2L1*. Part of the anti-apoptotic BCL-2 family.

BCLXL_T - BCLXL transcript

BAD- Bcl-2-associated death promoter, apoptosis sensitizer family representative

BIM- Bcl-2-like protein 11, apoptosis activator family representative

BIM_T – transcript of BCl-2-like protein 11

Apoptosis - Propensity of the cell to commit to programmed cell death.

Alpelisib – Isoform-specific drug inhibitor of PI3K alpha, also known as BYL719

Fulvestrant – drug inhibitor of the estrogen receptor, a type of selective estrogen receptor degrader (SERD)

Neratinib – drug inhibitor of EGFR and HER2

Palbociclib – drug inhibitor of CDK4/6

Everolimus) – mTORC1 and mTORC2 inhibitor

Trametinib – drug inhibitor of MAPK signaling (specifically, of MEK1 and MEK2)

Ipatasertib – drug inhibitor of AKT

S63845 - drug inhibitor of MCL1. A type of BH3 mimetic.

Navitoclax - drug inhibitor of BCL-2 and BCL-XL. A type of BH3 mimetic.

#### Attractors, initial conditions, and source nodes

The nodes of the model denote either the state of an intracellular entity (e.g. gene transcript, protein, signaling molecule), a biological outcome ($Apoptosis$, $Proliferation$, $ER\_transcription\_feedback\_1$, $ER\_transcription\_feedback\_2$), a drug ($Alpelisib$, $Fulvestrant$, $Neratinib$, $Palbociclib$, $Everolimus$, $Trametinib$, $Ipatasertib$, $S63845$, $Navitoclax$), or a cellular microenvironment ($HER3\_microenviroment$, $ER\_microenviroment$).

Nodes that are not regulated by other network nodes are called source nodes. They include all nodes denoting drugs and microenvironments, and also include 18 nodes that denote the basal transcriptional status of genes ($IGF1R\_T$, $HER2$, $HER3\_T$, $PDK1$, $SGK1\_T$, $PIM$, $PTEN\_T$, $p27\_T$, $p21\_T$, $BIM\_T$, $BCL2\_T$, $BCLXL\_T$, $MCL1\_T$, $ER$, $FOXA1$, $PBX1$, $MYC\_T$, and $Rb\_T$) . There are also some nodes that are only regulated by nodes denoting drugs ($CDK4/6$ and $mTORC2$) and which we will also refer to as source nodes.

The state of the majority of these source nodes is dictated by our context of interest: ER+ *PIK3CA* mutant breast cancer and the cell lines MCF7 and T47D. For this context and in both cell lines we have that 11 of these source nodes are active (i.e., have the node state 1; $IGF1R\_T$, $PTEN\_T$, $p27\_T$, $BIM\_T$, $MCL1\_T$, $ER$, $FOXA1$, $PBX1$, $MYC\_T$, $CDK4/6$, and $Rb\_T$) and 7 are inactive (i.e., have the node state 0; $PDK1$, $SGK1\_T$, $PIM$, $HER2$, $HER3\_T$, $mTORC2$, $BCL2\_T$). The state of $BCLXL\_T$ and $p21\_T$ is cell line-specific. The state of the source nodes denoting drugs and cellular microenvironments are set according to the treatment that is being modeled.

The model has 32 attractors (all of them steady states) under the selected source node states and under the inactive state for all drugs and cellular microenvironments: 4 cancerous states with high survivability ($Apoptosis=0$) and 28 cancerous states with low survivability ($Apoptosis=1,2,3$). Each of these steady states can either be primed for apoptosis ($BIM\_T=BIM=1$) or unprimed ($BIM\_T=BIM=0$), and can have the node $ER\_transcription\_feedback\_2$ (which denotes the memory of previous activation of the negative feedback loop from $ER\_transcription$ *to* $ESR1$) either active or inactive.

We use as initial state during simulations a mix of the two high survivability cancerous steady states with $ER\_transcription\_feedback\_2=0$, namely, the primed (with probability 1/3) and the unprimed state (with probability 2/3).

#### Update probability of the nodes

The update probability of node $k$ ($p_{k}$) depends on whether the node is categorized as a fast ($p_{k}=p_{fast}$) or a slow ($p_{k}=p_{slow}$) node. We categorize nodes into fast or slow based on whether activation of the node denotes a (fast) signaling event or a (slow) transcriptional or translational event. Following our previous model (Gómez Tejeda Zañudo, Scaltriti, and Albert 2017), the fast probability is chosen to be 5 times faster than the slow probability ($p_{fast}=5 p_{slow}$).

The fast nodes are: AKT, BAD, cycE_CDK2, cycD_CDK4/6, pRb, EIF4F, FOXO3, HER2/3, KMT2D, MAPK, mTORC1, mTORC2_pm, p21_p27, PDK1_pm, PI3K, PIM, PTEN, PIP3, PRAS40, RAS, S6K, SGK1, Translation, TSC, ER_microenvironment, HER3_microenvironment, Alpelisib, Fulvestrant, Neratinib, Palbociclib, Everolimus, Trametinib, Ipatasertib, S63845, Navitoclax

The slow nodes are: BIM, BIM_T, BCL2, BCL2_T, BCLXL, BCLXL_T, MCL1, MCL1_T, HER2, HER3, HER3_T, cycE_CDK2_T, IGF1R, IGF1R_T, ER, ESR1, FOXA1, PBX1, ER_transcription, cyclinD, CDK46, Rb_T, E2F, p27_T, p21_T, p21_p27_T, SGK1_T, PDK1, ER, mTORC2, FOXO3_Ub, Apoptosis, Proliferation, MYC, MYC_T, MYC_targets, ER_transcription_feedback_1, ER_transcription_feedback_2, PTEN_T.

#### Normalized value of Apoptosis and Proliferation ($Apoptosis_{norm}$ and $Proliferation_{norm}$)

The nodes $Apoptosis$ and $Proliferation$ denote the propensity of a cell to commit to programmed cell death and cell cycle progression, respectively. $Apoptosis$ and $Proliferation$ are multi-level nodes and each of them can take the discrete values 0, 1, 2, or 3. To facilitate the interpretation of the state of the nodes and weigh differently each of the states, we use the normalized variables $Apoptosis_{norm}$ and $Proliferation_{norm}$,

$$Apoptosis_{norm} = w_{A1}I(Apoptosis,1)+w_{A2}I(Apoptosis,2)+w_{A3}I(Apoptosis,3)$$

$$Proliferation_{norm} = w_{P1}I(Proliferation,1)+w_{P2}I(Proliferation,2)+w_{P3}I(Proliferation,3)$$

Here $I(v,s)$ is the indicator function ($I(v,s)=1$ if $v=s$, and $I(v,s)=0$ otherwise) and the normalized values are weighted exponentially (that is, $w_{Xi}=1/2^{3-i}$, with $X=A,P$and $i=1,2,3$). These weights are such that $Apoptosis=0, 1, 2, 3$ results in $Apoptosis_{norm}=0, 0.25, 0.5, 1$, respectively, and the same for $Proliferation$. This is the same principle we followed in the previous model, but the weights are modified for the case of Proliferation since the updated model has 4 node states for Proliferation, while the previous model had 5. The weight of $Apoptosis=1$ ($w_{A1}$) in $Apoptosis_{norm}$ is one of the differences in the cell line-specific models.

#### Cell line-specific models

The cell line-specific models for MCF7 and T47D use the updated network model. The MCF7-specific and T47D-specific models differ only in three aspects:

1. The source node $BCLXL\_T$, which denotes the expression of the gene encoding for BCL-XL, is active in T47D ($BCLXL\_T=1$) but not in MCF7 ($BCLXL\_T=1$), in agreement with Fig. 3E. This also results in a difference in the state of $BCLXL$ in the attractors of MCF7 ($BCLXL=0$) and T47D ($BCLXL=1$).
2. The source node $p21\_T$, which denotes the expression of the gene *CDKN1A*, is more active in MCF7 ($p21\_T=2$) than T47D ($p21\_T=1$), in agreement with Fig. 5E.
3. The weight of $Apoptosis=1$ ($w_{A1}$) in $Apoptosis_{norm}$ is larger in MCF7 ($w_{A1}=0.25$) than in T47D ($w_{A1}=0.125$). This is in agreement with Figs. 2D and 3E.

#### FOXO3, ER microenvironment, HER3 microenvironment

The updated model includes nodes that denote microenvironments: a microenvironment that induces ER activity when ESR1 is upregulated ($ER\_microenvironment$) and a microenvironment that induces HER2/HER3 activity when HER3 is upregulated ($HER3\_microenvironment$). These microenvironment nodes are included so that the model can be consistent with both our experimental results on FOXO3 (which upregulates ESR1 and HER3) and previous work on resistance mechanisms to PI3Kα inhibitors in these microenvironments.

In addition to its tumor suppressor effect, FOXO3 acts as an oncogene in the context of PI3K inhibition in breast cancer. Increased FOXO3 nuclear activity through PI3K/AKT inhibition in breast cancer results in feedback activation of receptor tyrosine kinases (e.g. HER3 or IGF1R, [(Chandarlapaty et al. 2011; Muranen et al. 2012; Kodack et al. 2017)]) and ER transcriptional activity [(Bosch et al. 2015)]. A consequence of the dual tumor suppressor and oncogene effect of FOXO3 is that, depending on their relative strength, a decrease in FOXO3 activity can result in reduced sensitivity to PI3Kα inhibition (if the tumor suppressor effect dominates) or increased sensitivity (if the oncogenic feedback activation effect dominates). In the network model we previously built (Gómez Tejeda Zañudo, Scaltriti, and Albert 2017), $FOXO3=0$ led to a multi-faceted change in the response to alpelisib compared to the normal (unperturbed) response to alpelisib: reduction in apoptosis ($Apoptosis_{norm} =0.33$ vs $Apoptosis_{norm} =0.70$, through FOXO3’s effect on BIM) and also a reduction in proliferation ($Proliferation_{norm}=0.13$ vs $Proliferation_{norm} =0.25$, through FOXO3’s feedback activation effect on ESR1).

Given the reduction in alpelisib-induced apoptosis caused by FOXO3=OFF, we considered downregulation of FOXO3 as a potential resistance mechanism to alpelisib. However, since $FOXO3=0$ also resulted in a reduction in proliferation (and survivability) through feedback regulation of ER transcription, downregulation of FOXO3 could increase sensitivity to alpelisib in microenvironments in which the feedback activation effects dominate. For example, in an estrogen-rich microenvironment, reduced ER transcriptional activity caused by downregulation of FOXO3 might result in an increased sensitivity to alpelisib, and similarly in a HER3-activating environment (e.g. the brain microenvironment or a neuregulin-rich environment [(Kodack et al. 2017)]). Our previous model did not explicitly include the presence of these microenvironments, and implicitly assumed an ER microenvironment.

The logical rules of the microenvironment target nodes in the updated network model are such that $ER\_microenvironment$ is necessary for long-term $ESR1=2$ ($f_{ESR1\_2}=1$) and $HER3\_microenvironment$ is necessary for $HER2/3=2$ ($f_{HER2\_3\_2}=1$). The updated rules for these nodes are the same as the one in the original model except that each has an extra term:

$$f_{ESR1\_2}=f_{ESR1\_2}(previous model) and (ER\_microenvironment or not ER\_transcription\_feedback\_2)$$

$$f_{HER\_2\_3\_2}=f_{HER\_2\_3\_2}(previous model) and (HER3\_microenvironment)$$

The updated network model is able to recapitulate the reduced sensitivity to PI3Kα inhibitors in HER3 and ER microenvironments, for which FOXO3-mediated upregulation of HER3 and ESR1 has a greater weight on survival than the tumor suppressor role of FOXO3. The model also recapitulates the reduced PI3Kα inhibitor sensitivity to FOXO3 knockdown we observed in our experiments, in which the absence of HER3 and ER microenvironments leads to the tumor suppressor effect of FOXO3 to dominate. In particular, in the updated model FOXO3 knockdown results in both a reduction in apoptosis and an increase in proliferation in response to PI3Kα inhibitors.

#### Apoptosis

In the previous model, the rule for the node $Apoptosis$ assumed that alpelisib alone would induce cell death. This was because alpelisib resulted in the transcriptional upregulation of BIM, the phosphorylation and activation of BAD, and the transcriptional downregulation of MCL1. The previous model also assumed that the combination of alpelisib and fulvestrant would result in an increase in cell death compared to either of them alone (through upregulation of BCL2). The regulatory function for $Apoptosis$ also assumed that downregulation of MCL1 or BCL2 alone would not induce cell death. In our experiments, alpelisib did not result in a significant induction of cell death, and only showed a significant induction when combined with BH3 mimetics (Figs. 2D-E, 3C-D). Fulvestrant did not result in an increase in cell death, either alone or when combined with alpelisib. We did not observe a significant decrease in MCL1 protein expression in response to alpelisib and found that MCL1 inhibitor s63845 alone (or in combination with BCL-XL/BCL-2 inhibitor navitoclax) resulted in the induction of cell death (Figs. 2D-E, 3C-D).

To capture our experimental results, we modify the regulatory function for $Apoptosis$ so that alpelisib alone (through its induction of BIM and BAD) will only have a small effect on apoptosis ($Apoptosis=1$). For this to be the case, we also needed to remove the edge responsible for the downregulation of MCL1 caused by alpelisib, and instead have MCL1 depend only on the basal transcriptional status of MCL1 ($MCL1\_T$) and the presence of the drug s63845. We also removed the edge responsible for the induction of cell death by fulvestrant (from $ER\_transcription$ to $BCL2$) since we did not observe a significant induction of apoptosis by fulvestrant. Since BH3 mimetics alone had a larger effect on apoptosis than alpelisib alone, we modified the rule so that low activity of MCL1, BCL-XL, and BCL2 resulted in a larger effect on apoptosis ($Apoptosis=2$). The largest effect on apoptosis ($Apoptosis=3$) was the combination of low activity of MCL1, BCL-XL, and BCL2, and induction of BIM.

$$f_{Apoptosis}=(BIM and BAD) or Apoptosis$$

$$f_{Apoptosis\_2}=not (MCL1 or BCL2 or BCLXL) or Apoptosis\_2$$

$$f_{Apoptosis\_3}=(BIM and not (MCL1 or BCL2 or BCLXL)) or Apoptosis\_3$$

$$f_{MCL1}=MCL1\_T and not S63845$$

$$f_{BCLXL}=BCLXL\_T and not Navitoclax$$

$$f_{BCL2}=BCL2\_T and not Navitoclax$$

#### Negative interaction from ER transcription to ESR1

Alpelisib-induced upregulation of ESR1 is only transient (Bosch et al. 2015; Toska et al. 2017). It is unclear what the mechanism behind ESR1’s upregulation being transient is, but we enforce it through a negative feedback loop from $ER\_transcription$ *to* $ESR1$. In particular, we use nodes ($ER\_transcription\_feedback$, $ER\_transcription\_feedback\_2$) to encode time delays and make this negative feedback loop slow.

$$f_{ER\_transcription\_2}=f_{ER\_transcription\_2} (previous model) =KMT2D and FOXA1 and PBX1 and ESR1\_2$$

$$f_{ER\_transcription\_feedback}=ER\_transcription\_2$$

$$f_{ER\_transcription\_feedback\_2}=ER\_transcription\_feedback or ER\_transcription\_feedback\_2$$

$$f_{ESR1\_2}=f_{ESR1\_2}(previous model) and (ER\_microenvironment or not ER\_transcription\_feedback\_2)$$

#### MYC targets

In the previous model, the node denoting cap-dependent translation ($Translation$), which is regulated by the mTORC1 pathway, was a direct regulator of the state of $Proliferation$. This was based on previous work on the importance of cap-dependent translation in response to PI3K/mTOR inhibitors (Muranen et al. 2012) and the biological knowledge on mTOR-mediated translational control (Ma and Blenis 2009). Recent work on PI3K inhibitors, PI3K pathway signaling and MYC-driven cancer made us reconsider the direct interaction from $Translation$ to $Proliferation$, and we instead assume that this interaction is mediated by a new node, $MYC\_targets$.

Very recent work on PI3Kα inhibitors found that multiple MYC target gene expression signatures (including the hallmark signatures (Liberzon et al. 2015)) are strongly downregulated in response to alpelisib in MCF7 and T47D cells, and that MYC overexpression can cause resistance to PI3Kα inhibitors (Donnella et al. 2018). This is consistent with past work showing that overexpression of MYC or of eIF4F (which is part of the mTORC1 pathway and a direct regulator of $Translation$ in our models) induces resistance to PI3K/mTOR inhibitors (Ilic et al. 2011; Muellner et al. 2011). These results are also consistent with very recent work finding a mutual exclusivity of *PIK3CA* and MYC alterations (Schaub et al. 2018), and the correlation of MYC and PI3K pathways signatures in breast and other cancers (Y. Zhang et al. 2017).

This work points to an overlap between the downstream effect of MYC and the mTORC1 pathway that is not fully captured in the previous model. To capture this overlap, the updated model includes the new node $MYC\_targets$ (the name of the MYC hallmark signatures in (Liberzon et al. 2015)), which denotes genes regulated by MYC and by mTORC1-dependent translation through the EIF4 factors.

The regulatory function for $MYC\_targets$ is such that either Translation and basal MYC activity ($MYC=1$) or high MYC activity alone ($MYC=2$) can result in high activity of MYC targets ($MYC\_targets=2$). Translation or MYC activity ($MYC\geq1$) alone can result in in intermediate activity of MYC targets ($MYC\_targets=1$).

$$f_{MYC\_targets}=(MYC\_2 or MYC) or Translation$$

$$f_{MYC\_targets\_2}=(MYC\_2) or (MYC and Translation)$$

#### Proliferation, pRb, and E2F

In the updated model, $pRb$, $E2F$, and $Proliferation$ each have one less state compared to the previous model (in the updated model $pRb$ and $E2F$ have 3 states and $Proliferation$ has 4 states). This change is motivated by the relatively small effect on PI3Kα inhibitor sensitivity seen by FOXO3 and RB1 knockdown compared to what was seen for other PI3Kα inhibitor resistance mechanisms (see e.g. (Le et al. 2016)).

In the previous model, $pRb$ and $E2F$ could have 4 states, which reflected distinct degrees of phosphorylation of Rb and their distinct ability to bind and inhibit the E2F family of transcription factors: unphosphorylated, multiple hypo-phosphorylated states (e.g. phosphorylation at 1 or 2 sites), and hyper-phosphorylated (Bertoli and de Bruin 2014; Lundberg and Weinberg 1998; Ezhevsky et al. 1997). The unphosphorylated Rb binds and inhibits the E2F family, while the ability to inhibit E2F is lower in the hypo-phosphorylated forms, and the lowest in the hyper-phosphorylated form.

Given that our experiments showed that RB1 knockdown has a small effect on sensitivity to alpelisib (Fig. 6), this suggests that we need less than 4 states of Rb to capture its effect on its targets, $E2F$ and $Proliferation$. The number of states of $pRb$ may not be fully dictated by the phosphorylation states of Rb. Since $Proliferation$ denotes the propensity of cells to enter the cell cycle, a more appropriate interpretation is to consider $pRb$ as the activity level of Rb in terms of its inhibitory effect on E2F and on the proliferation propensity (where $pRb=2$ denotes no anti-proliferative effect and $pRb=0$ is the maximum anti-proliferative effect) .

Using the above interpretation, the activity levels of $pRb$ would still depend on the activity of the cyclin E and cyclin D complexes (nodes $cyclinE/CDK2$ and $cycD/CDK4/6$), which regulate the activation of Rb during cell cycle progression. We assume that $pRb=2$ requires the activity of both the cyclin D and E complexes ($cyclinE/CDK2=1$ and $cycD/CDK4/6=1$) or high activity of the cyclin D complex ($cycD/CDK4/6=2$). For $pRb=1$, the activity of either complex is sufficient. In addition, since the loss of Rb results in no inhibitory effect of Rb on E2F family members, we encode this in the rules by enforcing that $Rb\_T=0$ results in $pRb=2$ (i.e., no inhibitory effect on the node $E2F$).

$$f_{pRb}=(cycD\_CDK46\_2 or cycD\_CDK46) or not Rb\_T$$

$$f_{pRb\_2}=((cycD\_CDK46 and cycE\_CDK2) or cycD\_CDK46\_2) or not Rb\_T$$

$$f_{E2F}=pRb$$

$$f_{E2F\_2}=pRb\_2$$

In the previous model, the number of states of the node $Proliferation$ (5 states) was determined by the number of states of its regulators $E2F$ (4 states) and $Translation$ (2 states), and the ability to reproduce previous knowledge on the response and resistance to PI3Kα inhibitors and other drugs in ER+ breast cancer. In the updated model, the number of states for the node Proliferation is constrained by our experimental results.

We assume that, in the absence of drugs, the cell has the highest proliferation propensity ($Proliferation=3$). With alpelisib, there is a much lower proliferation propensity. This is nevertheless not the lowest possible proliferation propensity, as the fact that combination of alpelisib and fulvestrant further reduces cell viability but not apoptosis suggests that proliferation propensity is even lower in this case. The lower proliferation propensity in response to alpelisib would then correspond to $Proliferation=1$, and the lowest one (e.g., in response to alpelisib and fulvestrant) would correspond to $Proliferation=0$. Resistance mechanisms to alpelisib such as knockdown of PTEN, knockdown of TSC1/2, or PIM1 overexpression, should allow the cell to stay at the highest proliferation propensity ($Proliferation=3$), while the reduced sensitivity seen in FOXO3 or RB1 knockdown in our experiments should result in a small decrease of proliferation propensity ($Proliferation=2$). To accomplish this ordering of states, we assume that the highest state of Proliferation ($Proliferation=3$) can be achieved only with the highest activity of MYC targets ($MYC\_targets=2$) and an active state of E2F ($E2F\geq1$). The intermediate state ($Proliferation=2$, corresponding to the reduced alpelisib sensitivity seen for knockdown of FOXO3 or RB1) can be achieved by high activity of E2F ($E2F= 2$). The lowest active state of $Proliferation$ ($Proliferation=1$) can be achieved with the activity of either MYC targets ($MYC\_targets\geq1$) or E2F ($E2F\geq1$)

$$f_{Proliferation}=MYC\_targets or E2F$$

$$f_{Proliferation\_2}=E2F\_2$$

$$f_{Proliferation\_3}=MYC\_targets\_2 and (E2F\_2 or E2F)$$
