## Supplemental Figures for "Cell line-specific network models of ER+ breast cancer identify PI3Kα inhibitor sensitivity factors and drug combinations"

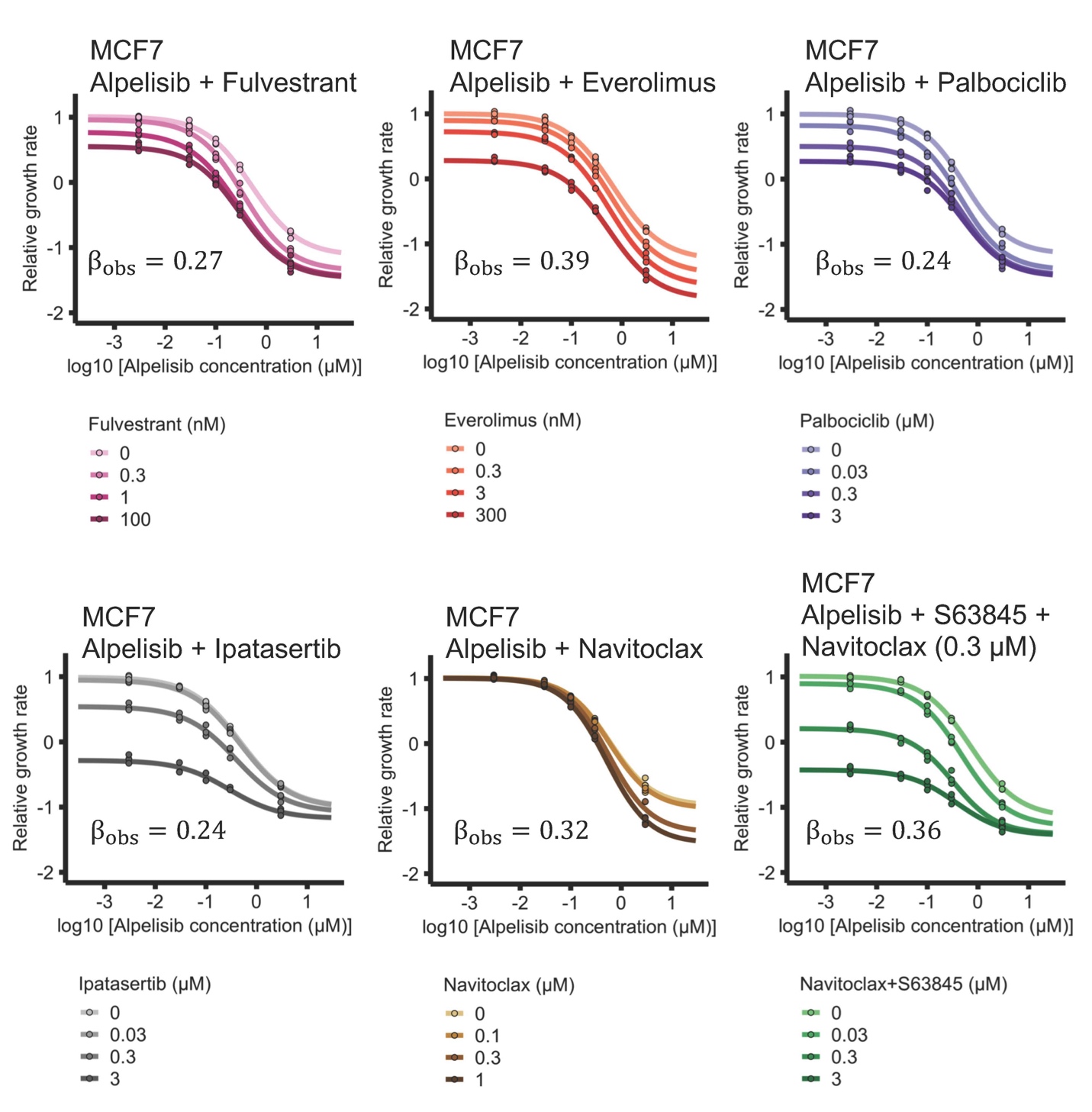

**Supp. Fig. S1. Combination and synergistic efficacy of alpelisib with selected clinically relevant drugs and BH3 mimetics in MCF7.** The selected clinically relevant drugs are fulvestrant (a selective estrogen receptor degrader), everolimus (an inhibitor of mTOR), palbociclib (a CDK4/6 inhibitor), and ipatasertib (a pan-AKT inhibitor). The BH3 mimetics are navitoclax (a BCL-XL/BLC-2 inhibitor) and s63845 (a MCL1 inhibitor). Efficacy scores $\beta_{obs}$ are calculated using MuSyC (Meyer et al. 2019), and are such that $\beta_{obs}>0$ means synergistic behavior and $\beta_{obs}<0$ means antagonistic behavior.

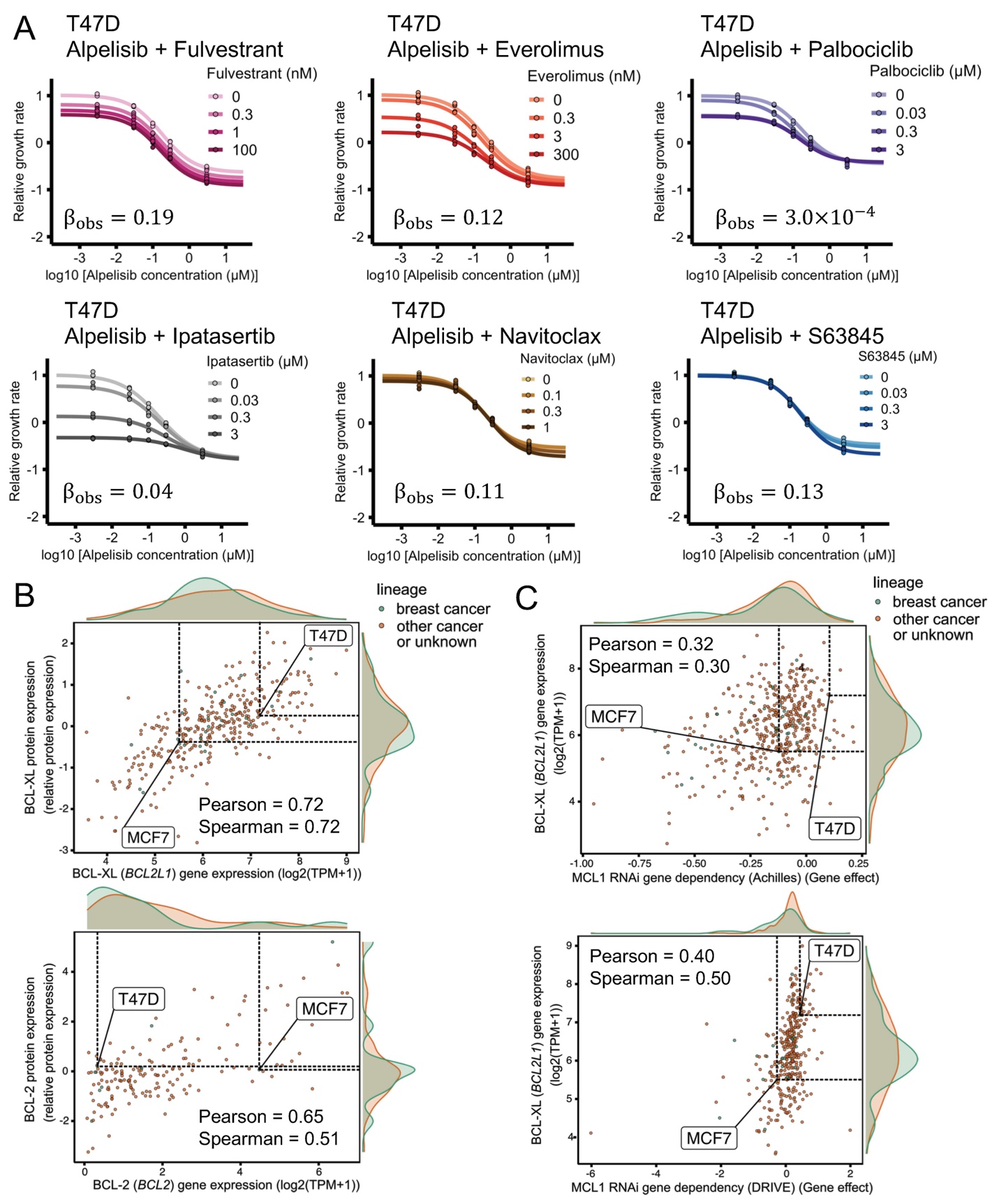

**Supp. Fig. S2. Combination of alpelisib with selected clinically relevant drugs and BH3 mimetics in T47D and gene/protein expression correlates for sensitivity to MCL1 knockout.** (A) Drug response curves of alpelisib combined with clinically relevant drugs and BH3 mimetics in T47D. The selected clinically relevant drugs are fulvestrant (a selective estrogen receptor degrader), everolimus (an inhibitor of mTOR), palbociclib (a CDK4/6 inhibitor), and ipatasertib (a pan-AKT inhibitor). The BH3 mimetics are navitoclax (a BCL-XL/BLC-2 inhibitor) and s63845 (a MCL1 inhibitor). Efficacy scores $\beta_{obs}$ are calculated using MuSyC (Meyer et al. 2019), and are such that $\beta_{obs}>0$ means synergistic behavior and $\beta_{obs}<0$ means antagonistic behavior. (B) Gene and protein expression of BCL-XL (*BCL2L1*) (top) and BCL-2 (*BCL2*) (bottom). The higher gene and protein expression of BCL-XL in T47D compared to MCF7 is consistent with T47D requiring navitoclax to be sensitive to combined alpelisib and s63845. (C) Sensitivity to MCL1 knockout in RNAi loss-of-functions screen datasets is strongly correlated with BCL-XL gene expression. MCF7 is more sensitive to MCL1 RNAi knockout than T47D (MCF7’s MCL1 gene effect is less than that of T47D), consistent with their differential sensitivity to s63845. Achilles dataset from Broad (top), DRIVE dataset from Novartis (bottom).

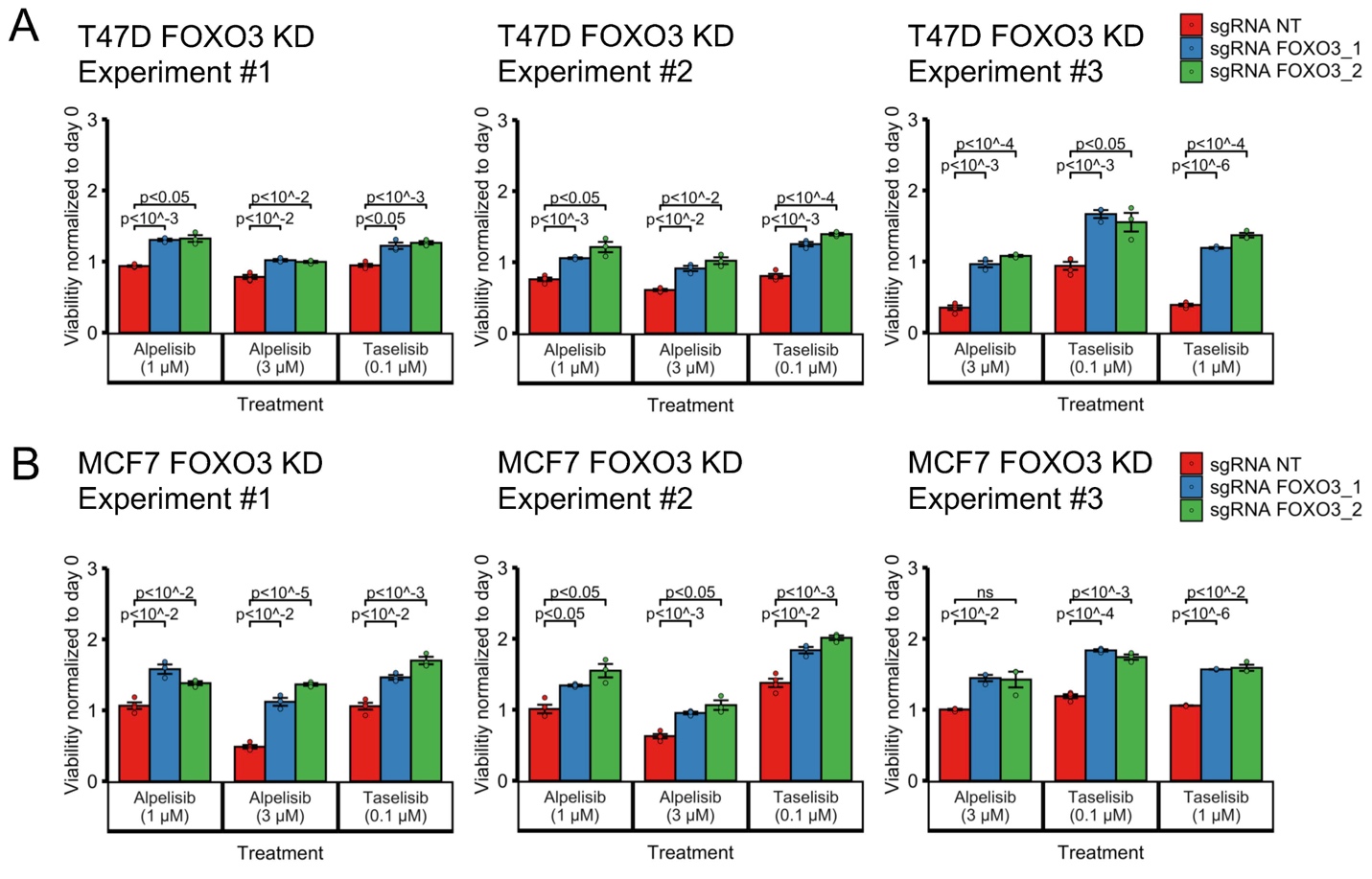

**Supp. Fig. S3. Reproducibility of FOXO3 knockdown’s reduced sensitivity to PI3Kα inhibitors alpelisib and taselisib.** Each experiment was done in a separate 96-well plates. Experiment #1 and #2 were done in the same week and are normalized according to the same day 0 untreated 96-well plate.

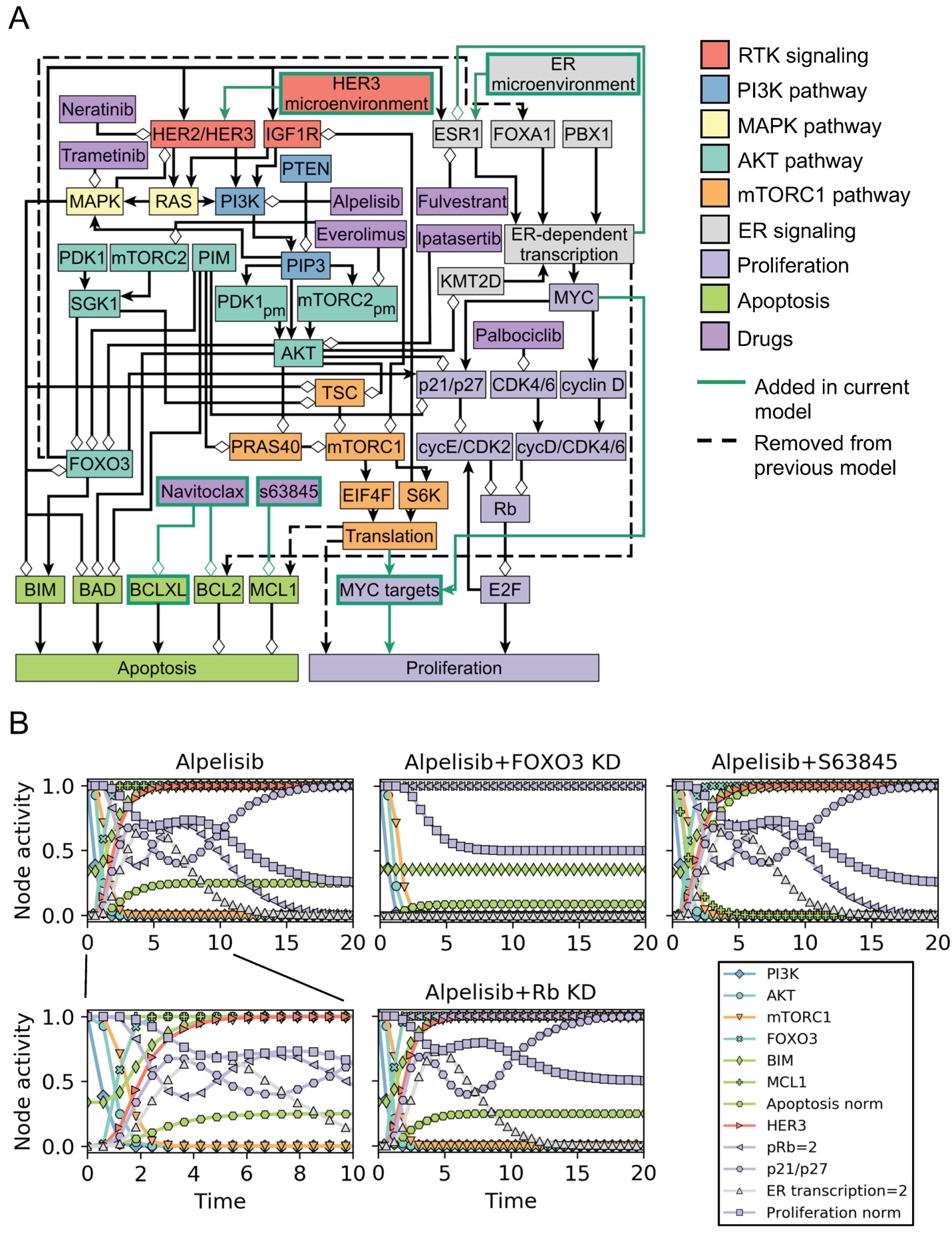

**Supp. Fig. S4. Updated network model of oncogenic signal transduction in ER+, *PIK3CA* mutant breast cancer.** (A) Cell line-specific models are based on the network model shown, which is an updated version of the network model of Zanudo et al. 2017. The updated network model incorporates new knowledge on ER+ breast cancer drug resistance (references (Hopkins et al. 2018; Donnella et al. 2018; Wander et al. 2020)) and the discrepancies between our experimental results and the Zanudo et al. 2017 model. The updated model has only slight modifications to the network structure compared to the previous model, and these are shown in panel A with a green node outline (for node additions), green arrow (for edge additions), or a dashed line (for edge deletions). More details on the modifications done to the model and the reasoning behind them can be found in the Supplemental Text and Supplemental File 1. (B) Timecourse of the node activity (average node state) in response to alpelisib in the updated network model. In particular, we show the timecourse for the MCF7-specific network model. In response to alpelisib, there model shows a quick inactivation of multiple pathways (AKT, MAPK, mTORC1), followed by the activation of $FOXO3$. Activation of $FOXO3$ results in the activation of $BIM$ (through transcriptional upregulation), which together with the inactivation of $BAD$ by $AKT$ results in an increase in $Apoptosis$. Activation of $FOXO3$ also activates $HER3$ and $ER transcription=2$ (through transcriptional upregulation of HER3 and ESR1), the latter of which is transient due to a negative feedback loop. The transient activation of results in the non-monotonic behavior of $Proliferation_{norm}$, which decreases because of the inactivation of MYC targets by the mTORC1 pathway and the activation of $p21/p27$ and $pRb=2$, but is transiently counteracted by $ER transcription=2$ . The timecourse for the response to alpelisib + FOXO3 knockdown shows how the response of multiple elements in the model depends on $FOXO3$ activation, including $BIM$, $Apoptosis_{norm}$, $p21/p27$, $HER3$, and the non-monotonic response of $pRb=2$, $ER transcription=2$, and $Proliferation_{norm}$. The timecourse for the response to alpelisib + s63845 differ to that of alpelisib alone in the state of the nodes $MCL1$ and $Apoptosis_{norm}$. The timecourse for the response to alpelisib + Rb knockdown differs to that of alpelisib alone only in the state of the nodes of $pRb$, $E2F$, and $Proliferation_{norm}.$

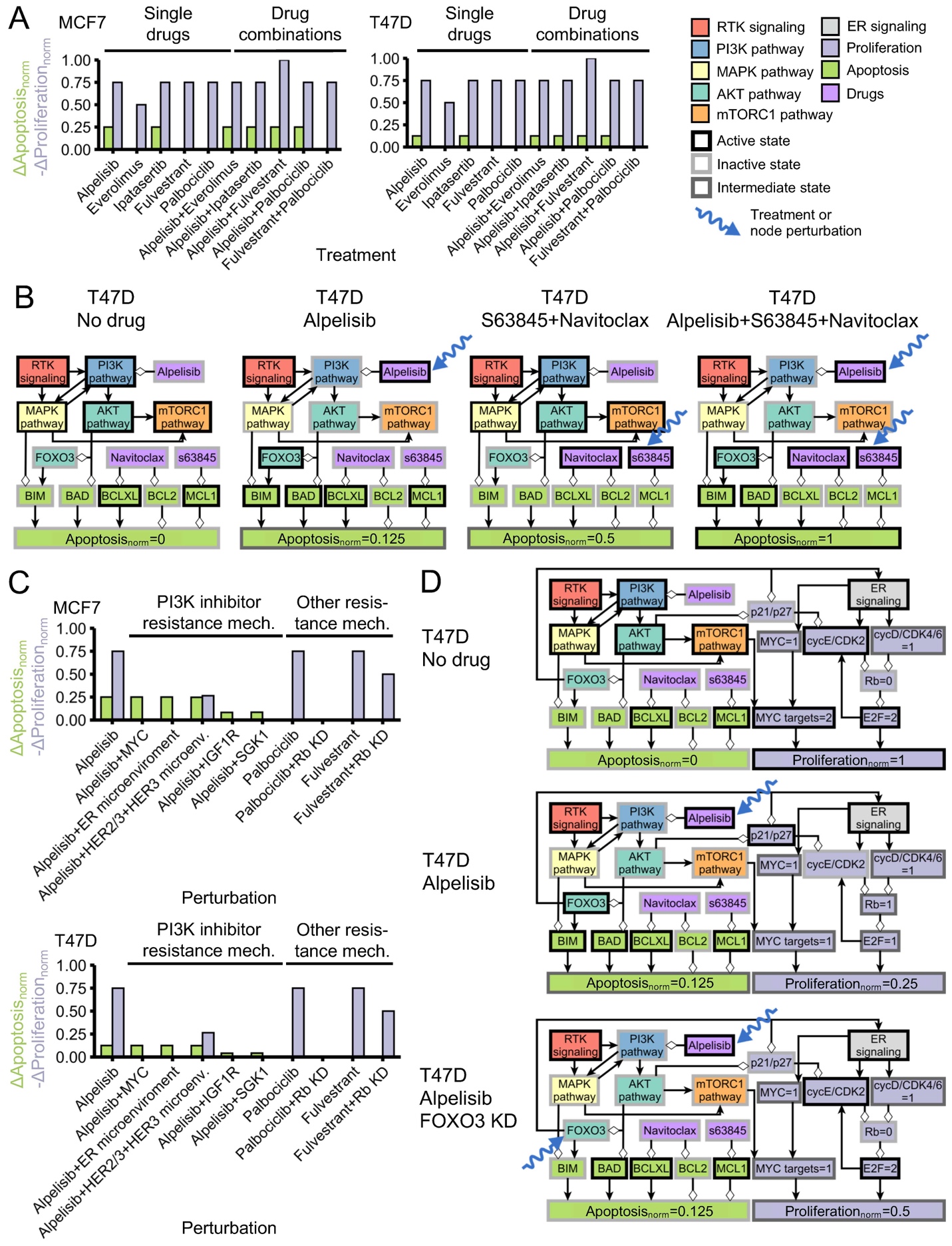

**Supp. Fig. S5. Additional drug combinations and resistance mechanisms in cell line-specific network models.** (A) Response of MCF7-specific and T47D-specific network modes to single drugs relevant in the context of ER+ breast cancer and the combination of these drugs with alpelisib. (B) T47D-specific model reproduces the observed cell death response to BH3 mimetics, alpelisib, and their combination seen in this cell line (compare to the experimental results in Fig. 3C). We show the state of the nodes that influence $Apoptosis$ in response to alpelisib, BH3 mimetics, and their combination. (D) Cell line-specific network models reproduce the drug resistance effect of PI3Kα inhibitor resistance mechanisms and other resistance mechanisms. Simulations in which a drug resistance mechanism is active show an increased survivability (reduced $Proliferation$ and/or increased $Apoptosis$) compared to the case of the drug alone (panel C). (E) T47D-specific model reproduces the reduced sensitivity to alpelisib caused by FOXO3 knockdown. We show the state of the nodes that influence $Apoptosis$ and $Proliferation$ in response to alpelisib, FOXO3 knockdown, and their combination. The models encodes the biological outcomes of cell death and proliferation using $Apoptosis_{norm}$ and $Proliferation_{norm}$, respectively, which are a weighted and normalized (between 0 and 1) measure of the state of the nodes $Apoptosis$ and $Proliferation$. Starting from a cancerous state of each model, we perform 10,000 simulations in which the specified treatment or perturbation is maintained throughout the simulation, and obtain the average value of $Apoptosis_{norm}$ and $Proliferation_{norm}$ at the end of the simulations. $\Delta Apoptosis_{norm}$ and $\Delta Proliferation_{norm}$ of a perturbation denote the difference with respect to the case of no perturbation ($Apoptosis_{norm}=0$, $Proliferation_{norm}=1$), and are such that decreased survivability means an increase in $\Delta Apoptosis_{norm}$ or $-\Delta Proliferation_{norm}$.

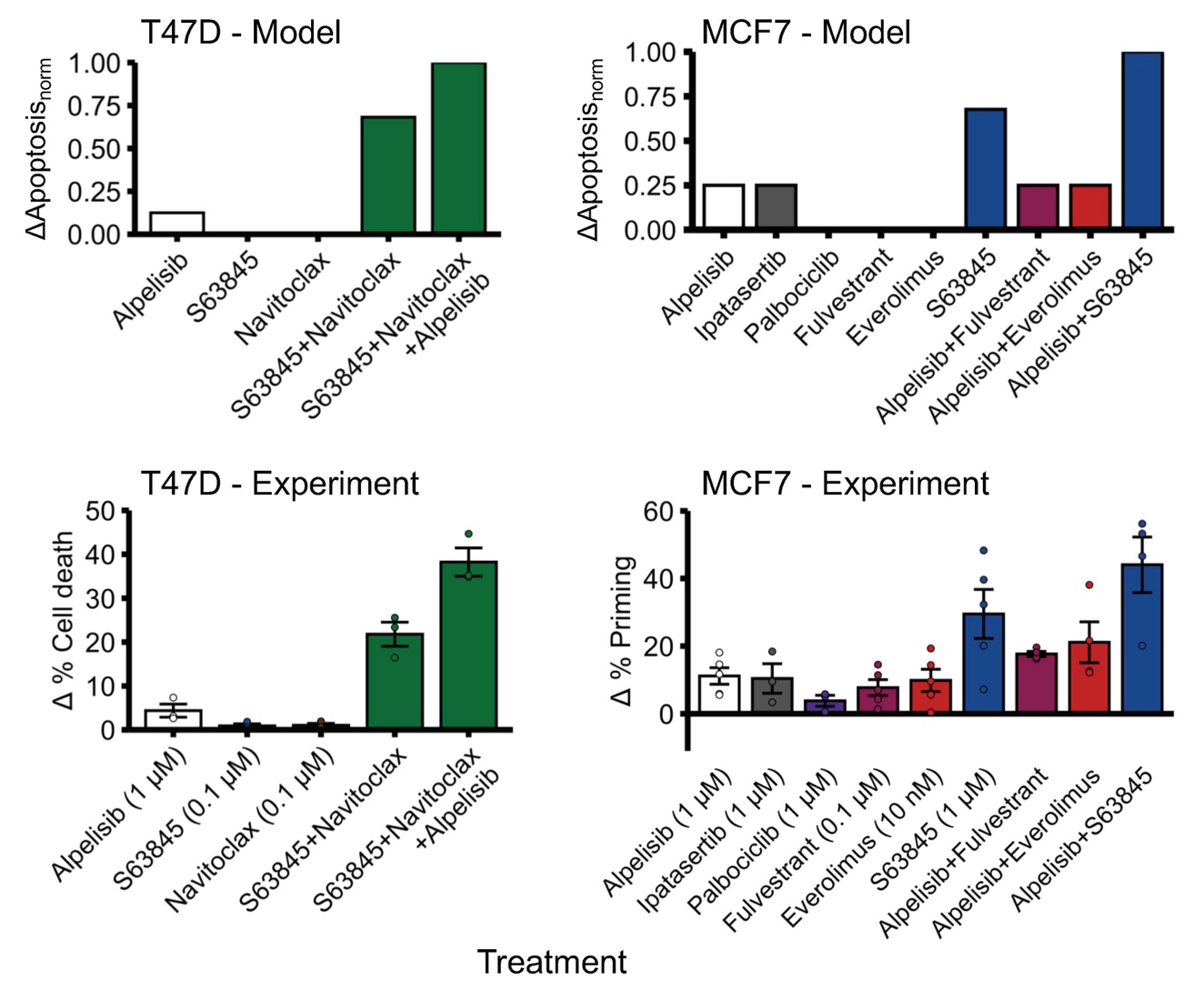

**Supp. Fig. S6. Cell line-specific network models reproduce the observed cell death response to alpelisib, BH3 mimetics, and other drugs.** The cell death response ($\Delta Apoptosis_{norm}$) in MCF7-specific and T47D-specific network models (top panel) is qualitatively similar to the experimentally observed cell death response in these cell lines (bottom panel). The experimental cell death results shown in the bottom panel are the same as in Figs. 2E and 3C. The models encode the biological outcomes of cell death using $Apoptosis_{norm}$, which is a weighted and normalized (between 0 and 1) measure of the state of the node $Apoptosis$. Starting from a cancerous state of each model, we perform 10,000 simulations in which the specified treatment or perturbation is maintained throughout the simulation, and obtain the average value of $Apoptosis_{norm}$ at the end of the simulations. $\Delta Apoptosis_{norm}$ of a perturbation denotes the difference with respect to the case of no perturbation ($Apoptosis_{norm}=0$).
